# c-MAF dependent perivascular macrophages regulate diet induced metabolic syndrome

**DOI:** 10.1101/2021.02.07.430147

**Authors:** Hernandez Moura Silva, Jamil Zola Kitoko, Camila Pereira Queiroz, Lina Kroehling, Fanny Matheis, Katharine Lu Yang, Christine Ren-Fielding, Dan Rudolph Littman, Marcelo Torres Bozza, Daniel Mucida, Juan José Lafaille

## Abstract

Macrophages are an essential part of tissue development and physiology. Perivascular macrophages have been described in tissues and appear to play a role in development and disease processes, although it remains unclear what are the key features of these cells. Here, we identify a subpopulation of perivascular macrophages in several organs, characterized by their dependence on the transcription factor c-MAF, displaying non-conventional macrophage markers including LYVE1, Folate receptor 2 and CD38. Conditional deletion of c-MAF in macrophage lineages caused ablation of perivascular macrophages in the brain and altered muscularis macrophages program in the intestine. In the white adipose tissue (WAT), c-MAF deficient perivascular macrophages displayed an altered gene expression profile, which was linked to an increased vascular branching into the tissue. Upon feeding on high fat diet (HFD), mice with c-MAF deficient macrophages showed improved metabolic parameters compared to wild-type mice, including less weight gain, greater glucose tolerance and reduced inflammatory cell profile in WAT. These results define c-MAF as a central regulator of perivascular macrophages cell identity and transcriptional program *in vivo* and reveal a novel role for this tissue resident macrophage population in the regulation of metabolic syndrome.

## INTRODUCTION

Since the seminal discovery of macrophages by Eli Metchnikoff(*1*), this immune cell type has been designated as a central player in innate immune responses. In the past decades, different flavors of macrophages have been uncovered and their importance in the ontogeny and function of several organs has become clear. For example, osteoclasts are crucial for proper bone remodeling, microglia for synaptic pruning, and lung alveolar macrophages for clearance of surfactant(*2–7*). Among the diverse subpopulations, perivascular macrophages have been associated with the regulation of tissue physiology. These cells function as a first barrier for invading pathogens or for potentially harmful blood-born substances in the tissues(*8*). However, tools to selectively target *in vivo* perivascular macrophages have not been developed so far. Therefore, deciphering molecules that contribute to the development or function of this cell subpopulation is fundamental to better understand their role in the organism physiology.

The white adipose tissue (WAT) of healthy adult animals contains a large population of innate and adaptive immune cells, numerically dominated by macrophages. The proportions and numbers of these immune cell types is known to change widely in conditions such as diet-induced obesity, genetically-determined obesity, fasting, infections, and aging, among other situations(*9–17*). We previously characterized the epididymal WAT (eWAT) macrophage cell populations fed normal diet (ND) in adult B6 mice. The predominant populations under these conditions were perivascular macrophages, which we named Vasculature-Associated adipose tissue Macrophages (VAM) 1, and VAM2(*15*). As their name suggest, both VAM populations are in very close contact with blood vessels, allowing them to rapidly endocytose a diverse array of macromolecules present in the bloodstream(*6, 15, 18–22*). Here, we searched for key molecules that define cell identity and functional properties of perivascular macrophages found across different tissues, revealing the transcription factor c-MAF as a fundamental factor to establish embryonically derived perivascular macrophages cell identity and function *in vivo*.

## RESULTS

### The Adipose tissue displays a subpopulation of perivascular macrophages dependent of the transcription factor c-MAF

Our previous published data revealed that VAMs display several atypical surface markers for macrophages (CD38^+^ LYVE1^+^ Folate Receptor 2 (Folr2)^+^ CD206^HIGH^)(*15*) (Fig. S1A). VAMs were previously divided in VAM1 (MHCII^HIGH^TIM4^LOW^) and VAM2 (MHCII^LOW^TIM4^HIGH^)(*15*), both representing macrophages intimately associated to blood vessels (Fig. 1A and fig. S1A). In human white visceral adipose tissue, we also observed the presence of perivascular macrophages tightly associated with the endothelium, displaying some of the markers of the Lyve1^+^CD206^HIGH^ VAMs (Fig. 1B and fig. S1B). This close contact with endothelial cells is associated with a high endocytic capacity for blood-borne macromolecules (Fig. S1C). To define the functional relevance of VAMs, we aimed at characterizing transcription factors preferentially or exclusively expressed by VAMs, hence to establish genetic targeting tools.

**Fig. 1.**
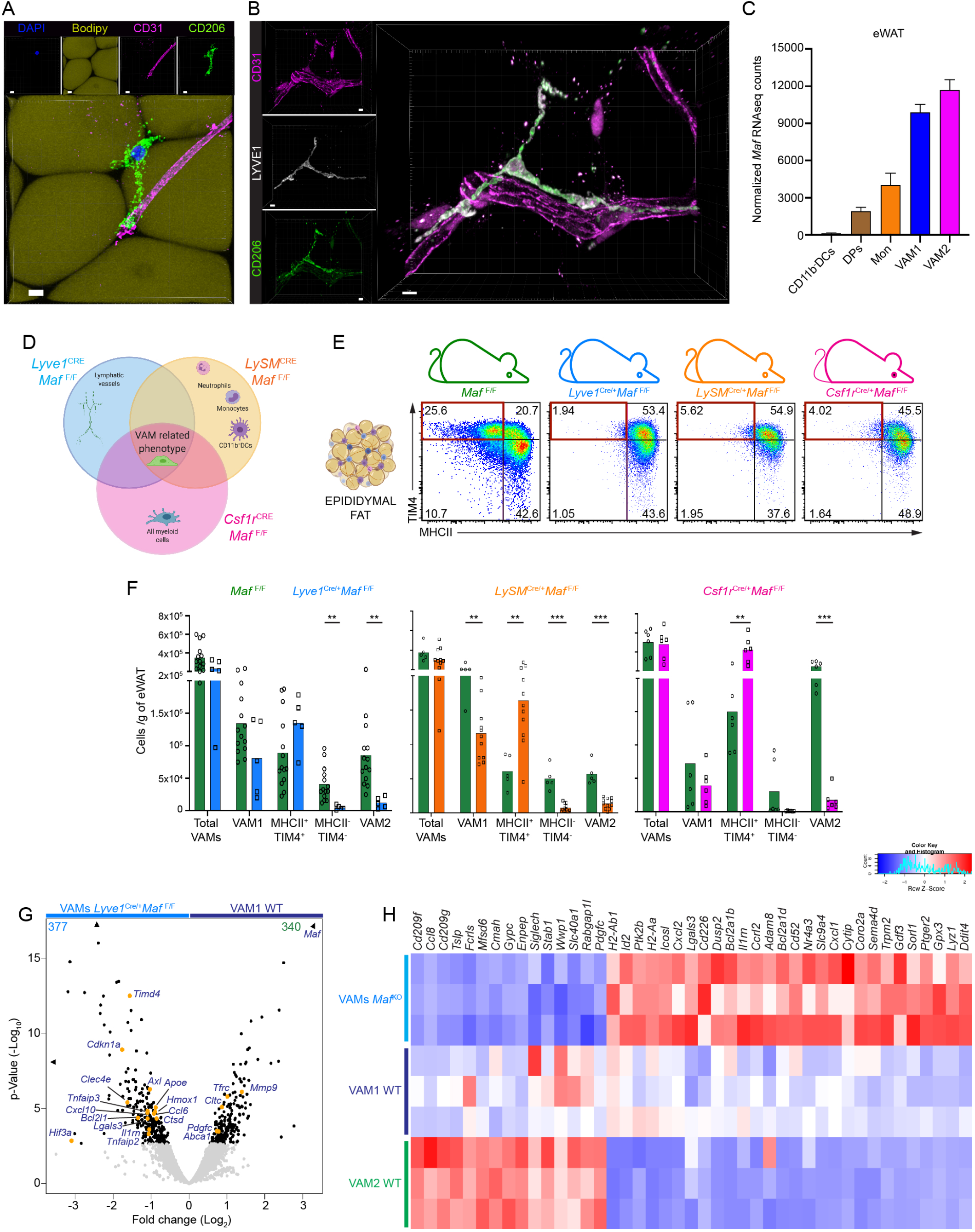
The visceral fat in mouse and humans harbors a *Maf*-dependent subpopulation of perivascular macrophages. (A) VAMs are closely associated with blood vessels in adult WT mice. eWAT full-mount sections stained with anti-CD206 (Green), anti-CD31 (Magenta), bodipy (yellow) and DAPI (blue). Scale Bars, 5 μm. 63x magnification. (B) Human visceral adipose tissue displays perivascular macrophages with a similar phenotype as described in mice. Confocal image of human clarified visceral adipose tissue stained with anti-CD206 (Green), anti-CD31 (Magenta) and anti-LYVE1 (white). Scale Bars, 15 μm. 63x magnification. Representative image. n=3. (C) *Maf* normalized RNAseq counts expressed by different subpopulations of myeloid cells in the eWAT as described previously(*15*). CD11b^-^DCs were gated as CD45^+^Lin^-^ CD11b^-^CD64^-^CD11c^+^MHCII^+^. n=3. (D) Schematic Venn diagram depicting the rationale of using 3 different cre drivers to ascertain the function of VAMs. (E) Impact of *Maf* ablation in VAMs macrophages in the eWAT. Each column represents VAMs from the animal depicted in the top. The gates in maroon color highlights the lost of cells with a typical VAM2 phenotype in the conditional knockout models used. Percentage of cells is shown. Representative dot plots of at least n=3. (F) Distribution of eWAT macrophages per gram of adipose tissue in the different conditional knockout models used in (E). Graph bar colors correspond to the models listed in (D, E). For each model, littermate controls were used as *Maf*^F/F^ WT controls. (G) Volcano plot comparing the top ≅8,000 genes expressed by VAM1 (MHCII^+^) from *Lyve1*^Cre^*Maf*^F/F^ and littermates controls from eWAT. Yellow dots highlight signature genes whose expression was modified by *Maf* gene deletion. Consolidated data from 3 animals in each group. Grey dots are genes non-significantly modulated. Numbers in the top corner are the total of differentially expressed genes. VAMs from both mice were purified by sorting as CD45^+^CD11b^+^CD64^+^Ly6C^-^ Ly6G^-^CD206^HIGH^MHCII^+^ cells. (H) *Maf* ablation leads to a transcriptional landscape distinct from VAM1 and VAM2 in the eWAT. Heat map of signature genes from *Maf* deficient. Each column represents one animal. The z-score of the gene expression profiles gives a scale to measure the differential expression. Each dot in the bar graphs represents one animal. Bar graphs display mean values. See also Fig. S1.

Analysis of our previously published epididymal WAT (eWAT) VAMs transcriptomics database (*15*) indicated that among all transcription factors sequenced, *Maf*, encoding the avian musculoaponeurotic fibrosarcoma oncogene homolog c-MAF, was highly expressed by VAMs (Fig. 1C). *Maf* expression in monocytes and CD11b^-^ DCs was much lower than in VAMs in the eWAT (Fig. 1C). *Maf* expression is also low in other tissue resident macrophages, such as microglia (Immgen.org and (*23*)), but observed in several non-hematopoietic cells, including Lymphatic Endothelial Cells (LEC). c-MAF was first reported to be expressed by myeloid cells in the 2000s when it was observed *in vitro* that it could modulated the expression of IL-10 in LPS stimulated murine macrophage cell lines (*24, 25*). More recently, it was show *in vitro* that c-MAF modulates the expression of several genes of M2-Like polarized bone-marrow derived macrophages (*26, 27*). c-MAF deficiency in mice leads to embryonic and postnatal lethality (*28, 29*). This has restricted the evaluation of c-MAF relevance for macrophages function *in vivo*. To circumvent that some groups tried to do bone-marrow chimeras by transferring fetal liver cells (E13-5) from *Maf* deficient embryos to lethally irradiated WT hosts (*30, 31*), a strategy that limits the observations mostly to radiosensitive/bone marrow derived cells. Therefore, it remains unclear how c-MAF contribute specifically to the function of low cycling resident perivascular macrophages such as VAMs *in vivo* (*15*). Thus, we first focused on the gene *Maf* as a target.

There is no available Cre driver to target specifically VAMs. *Lyve1*^Cre^ is also expressed by its original target, LEC (Immgen.org). *Lyz2* (LysM)^Cre^, the most commonly used Cre driver to target macrophages, is also expressed by other non-perivascular macrophages (except microglia) and is very highly expressed by granulocytes and monocytes, although LEC do not express *Lyz2*. Likewise, LEC do not express *Csf1r* (Immgen.org). *Csf1r*^Cre^ is expressed by yolk sac erythromyeloid progenitors (EMP), the precursors of tissue resident macrophages(*32, 33*), whereas *Lyz2* is not. Thus, we selected these three Cre drivers that share the property of being expressed by VAMs, while differing in their expression in cell types, to cross with *Maf*^Flox^ mice(*34*). We utilized a Venn diagram logic to use the three Cre drivers for complementary VAM targeting (Fig. 1D). All the conditional *Maf*-deficient mice survived into adulthood, gain weight normally, and have a normal gross anatomy (Data not shown). There was no obvious lymphatic drainage impairment in the *Lyve1*^Cre^*Maf*^F/F^ strain, which can be manifested by swollen extremities, due to liquid accumulation (Fig. S1D). Consistently, we observed a preserved capacity of the lymphatic vessels to drain evans blue dye from the foot pad to the popliteal lymph node (Fig. S1E, F). Furthermore, light-sheet microscopy using AdipoClear(*35*), which allows visualization of full architecture of the lymphatic vessels, showed that the eWAT lymphatic tree in *Lyve1*^Cre^*Maf*^F/F^ could be formed (Fig. S1G) and that the lymphatic vessels do not show any clear structural difference from those of WT mice (Fig. S1H).

VAMs were profoundly impacted by *Maf* gene deletion in the eWAT (Fig. 1E-H), observation recapitulated with all the Cre drivers, using flow cytometry, gene expression, and function. The population of VAM2 (Tim4^HIGH^MHCII^LOW^) was no longer observed, while VAM1 (Tim4^LOW^MHCII^HIGH^) was reduced by *Maf* targeting (Fig. 1E, F). These phenotypes were accompanied by a significant increase of cells in the normally underrepresented double-positive quadrant (Tim4^HIGH^MHCII^HIGH^) (Fig. 1E, F), resulting in an unaltered total number of VAMs (CD45^+^CD64^+^CD11b^+^CD206^HIGH^) among *Maf* mutants and WT littermates (Fig. 1F). Non-perivascular CD11c^+^CD64^+^ (DP) macrophages, which constitute a minor population in the eWAT from healthy mice, were unaffected both phenotypically and numerically by *Maf* gene deletion (Fig. S1I), consistent with a reduced (roughly 5X) expression of the *Maf* gene (Fig. 1C).

We subsequently analyzed the transcriptional profiles of eWAT VAMs upon *Maf* deletion. Compared to WT VAM1s, the gene expression pattern of *Maf* deficient VAMs had more than 700 differentially expressed genes, indicating that indeed *Maf* strongly shapes the transcriptional program of VAMs (Fig. 1G). In addition to the higher expression of *Timd4* compared to WT VAM1 cells, a number of anti-inflammatory genes, including *Il1rn, Tnfaip3, Hmox* and *Apoe*(*36*), were also upregulated in *Maf*-deficient VAMs (Fig. 1G, H). There was also upregulation of *Hif3a*/Nepas and *Cdkn1a* in *Maf*-deficient VAMs (Fig. 1G, H). *Hif3a* is a known repressor of the transcriptional activity of *Hif-1a* and *Hif-2a*/Epas(*37*), the two key genes that macrophages upregulate in response to hypoxia(*38*). *Cdkn1a* was shown to be instrumental in macrophage switching from a pro-inflammatory to an immunosuppressive state, and loss of *Cdkn1a* results in increased macrophage secretion of IFNβ and TNFα(*39*). There was reduced expression in *Maf*-deficient VAMs of *Ccl8* expression (Fig. 1H), a gene encoding a chemokine that, together with CCL2, attracts inflammatory monocytes(*40*) and is highly expressed by both VAM1 and 2 in the eWAT of WT mice(*15*). *Maf* ablation also resulted in downmodulation of *Pdgfc* by VAMs, a gene important to the cross talk between VAMs and the Csf-1-producing fibroblastic reticular cells, as well as a series of lectin and scavenger receptors, including as *Fcrls, CD209f, CD209g* and *Siglech* (Fig. 1H). Thus, the overall gene expression changes in VAM suggest that mice with conditional *Maf* deletion would be more resistant to adipose tissue inflammation, for example the one caused by high fat diet (HFD).

### *Maf* dependent VAM2s are predominantly embryonically sac derived

It is widely accepted that macrophages have two origins, one in yolk sac erythromyeloid progenitors (EMP), and the other in hematopoietic stem cells (HSC, fetal liver or bone marrow), via monocytes(*41–43*). To evaluate the origin of VAM1 and VAM2 we firstly carried out irradiated bone marrow (BM) chimeras (Fig. 2A). As expected, after two months all monocytes were donor-derived (Fig. 2B). However, a sizeable percentage of the eWAT VAMs (~30-50%) were host derived (Fig. 2B). VAM2s were only found in this group, retaining its TIM4^+^MHCII^LOW^ phenotype (Fig. 2C). Donor-derived macrophages were almost exclusively present in the VAM1 TIM4^-^ compartment (80-90%) (Fig. 2C). These experiments show that HSC do not contribute significantly to the VAM2 compartment. To independently confirm this data we performed kidney capsule transplant of neonatal eWAT to evaluate how the cell distribution would take place without irradiation (Fig. S2A). As observed in the BM chimeras, BM-derived macrophages were almost exclusively present in the VAM1 compartment while the few VAM2 found were reminiscent from the neonate donor (Fig. S2B). Finally, to verify a possible embryonic origin for VAM2 we used *CX3CR1*^CreER^ mice(*44*), as CX3CR1 is expressed by yolk sac EMPs(*41, 43*), but not HSCs. We carried out fate-mapping experiments by injecting a single low dose (to avoid abortion) of tamoxifen in a pregnant CX3CR1^CRE-ERT2^ x R26^-LSL-DsRed^ female mice (E8.5-9.5) (Fig. 2D). While the distribution, in adults, of VAM1 and VAM2 are approximately at a 50:50 percentage (Fig. 2E and(*15*)), the cells fate mapped at E9.5 were predominantly VAM2 (more than 70%, Fig. 2E). The above data indicates that VAM2 are embryonically derived while VAM1 are predominantly bone marrow derived.

**Fig. 2.**
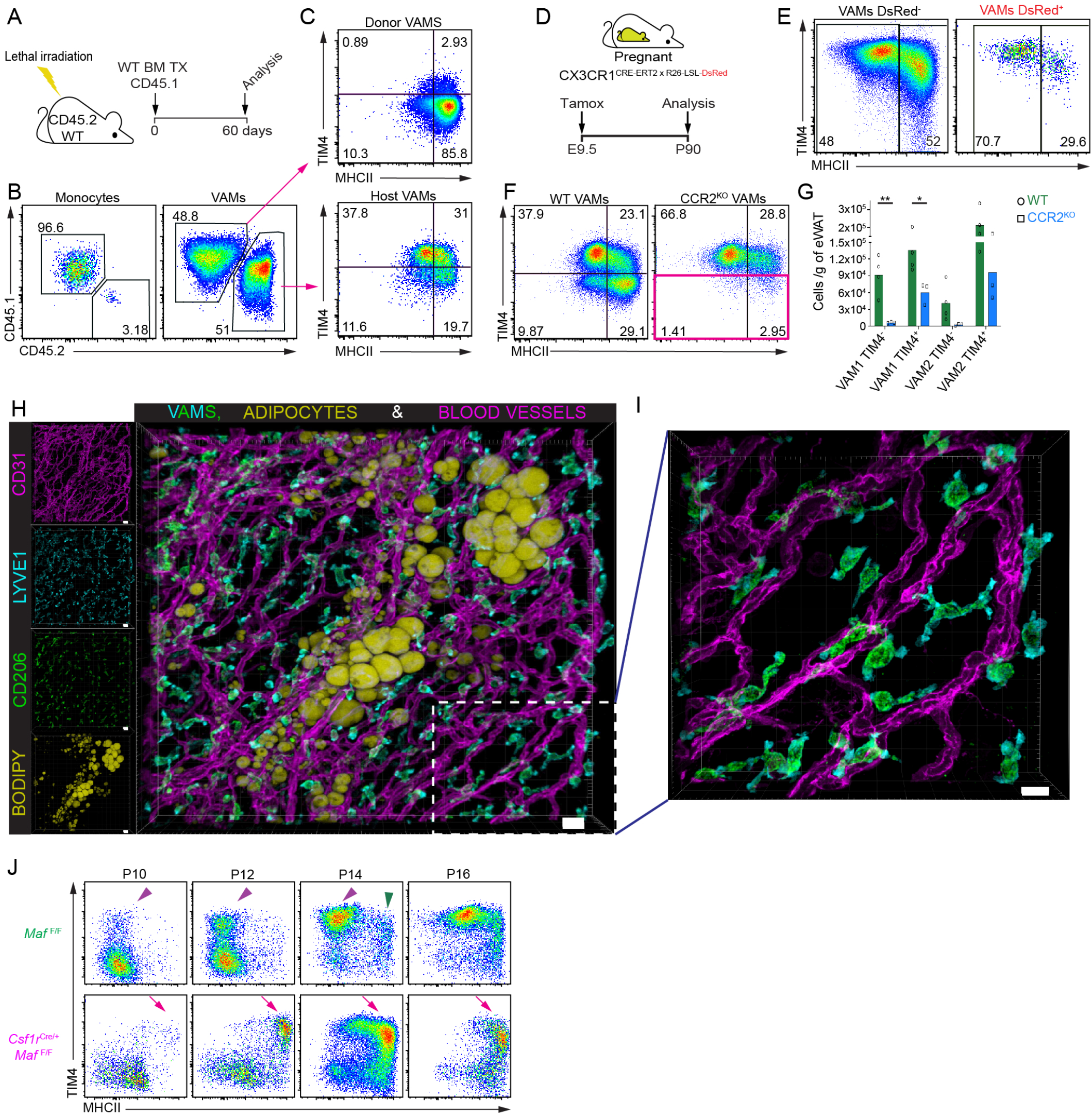
VAM2s are predominantly embryonically derived and show early dependence for *Maf* expression. (A-C) VAM2s can not be repopulated by bone marrow derived macrophages in bone marrow chimeras. (A) CD45.2 congenic WT mice were lethally irradiated and adoptively transferred bone marrow (BM) cells from a CD45.1 congenic donor. After 60 days of the BM transfer the eWAT was harvested and analyzed. (B) Distribution of host (CD45.2) and donor cells (CD45.1) among monocytes (gated as CD45^+^Lin^-^CD11b^+^CX3CR1^+^MHCII^-^Ly6C^HIGH^) and VAMs (gated as CD45^+^Lin^-^CD11b^+^CD64^+^CD206^HIGH^). Representative dot plot. n=4. Percentage of cells is shown. (C) Phenotype of host *versus* donor VAMs depicted in (B). Percentage of cells is shown. (D) Yolk sac fate mapping of EMPs. Pregnant *CX3CR1*^CRE-ERT2xR26-LSL-DsRed^ mice were treated with tamoxifen on day 9.5 of pregnancy to label EMPs with DsRed color. 90 days post birth the offspring was analyzed. (E) Distribution of VAMs from the animals depicted in (D). DsRed^+^ cells have an EMP origin. Representative dot plot. n=4. Percentage of cells is shown. (F), Analysis of VAMs from CCR2^KO^ mice. Representative dot plot of VAMs from CCR2^KO^ mice and littermate controls. Gate in pink highlight cell populations impacted by CCR2 deficiency. Percentage of cells is shown. (G), Absolute cell numbers of cells shown in (F). (H-I) Confocal microscopy of eWAT from P5 WT mice stained with anti-CD206 (Green), anti-CD31 (Magenta), anti-LYVE1 (Cyan) and Bodipy (Yellow). Scale Bars, 20 μm. 20x magnification. Images representative of at least n = 3. (I) High magnification of the quadrant depicted in (H). Scale Bars, 10 μm. 63x magnification. (J) eWAT MHCII^+^TIM4^+^ VAMs appear early on in the development of the eWAT in animals *Csf1r*^Cre^*Maf*^F/F^. Newborns were sacrificed in the ages depicted above each column and the rudimentary eWAT analyzed by flow cytometry. Purple arrowhead indicated the early appearance of VAM2 in WT mice at P10-P12. Green arrowhead shows VAM1 emergence at P14. Magenta arrows show the emergence of MHCII^+^TIM4^+^ VAMs by P10-P12 in *Csf1r*^Cre^*Maf*^F/F^ mice. Representative images. n=3. See also Fig. S2.

Several reports in the literature describe that the CC motif chemokine receptor 2 (CCR2) is a fundamental chemokine receptor for tissue homing of BM-derived monocytes that give rise to BM-derived macrophages(*45–47*). Hence, we decided to evaluate CCR2 knockout (KO) animals to verify the impact in the distribution of VAM1. When we examined the VAMs compartment in the eWAT, virtually all cells TIM4^-^ are vanished while the VAM2 are maintained in CCR2 deficient animals (Fig. 2F, H), further supporting their embryonically origin, while VAM1 are predominantly bone-marrow derived.

The dichotomy in the developmental origin between VAM1 and VAM2 led us to investigate how the populations are distributed during the establishment of a mature eWAT. First, we evaluated whether macrophages display adult morphological features in the developing fat pad. Confocal microscopy from P5 animals indicated that the developing eWAT resembles an empty sac with few developing adipocytes, an extensive blood network and VAMs with a different morphology (Fig. 2H). Instead of wrapping around blood vessels, a substantial fraction of the VAMs display a phenotype just emitting projections towards the blood vessels, far away from adipocytes (Fig. 2H, I). This environmental difference during the eWAT development reflected in the kinetics of VAM1 and VAM2 distribution in the organ. In WT pups, the first eWAT VAMs to appear, at P10, had a VAM2 phenotype, and were present in large numbers by P12; macrophages with a VAM1 phenotype only clearly appeared at day P14 (Fig. 2J, top row – purple and green arrowheads respectively).

Finally, these developmental differences among VAM1 and VAM2 led us to investigate whether the increase in Tim4^HIGH^MHCII^HIGH^ VAMs in *Maf* deficient mice (*see* Fig. 1E, F) was due to an early impact of *Maf* ablation in developing VAMs. In contrast to WT littermates, *Csf1r*^Cre^ *Maf*^F/F^ pups at P10 already displayed VAMs with an unusual Tim4^HIGH^MHCII^HIGH^ phenotype observed in adult conditional *Maf*-deficient mice (Fig. 2J, bottom row – arrows). These data point to an important role for c-MAF early in the establishment of VAM2s cell identity.

### VAM-like *Maf*-dependent perivascular macrophages are found in the brain and gut

To further understand if the CD38^+^ LYVE1^+^ FOLR2^+^ CD206^HIGH^ phenotype was a peculiarity of eWAT VAMs or a general feature of perivascular macrophages we investigated the brain, large and small intestine. Interestingly, we found prominent vascular-associated macrophages populations in the brain and gut that share the same markers (Fig. 3A-C, fig. S3A-C and fig. S4A-C).

**Fig. 3.**
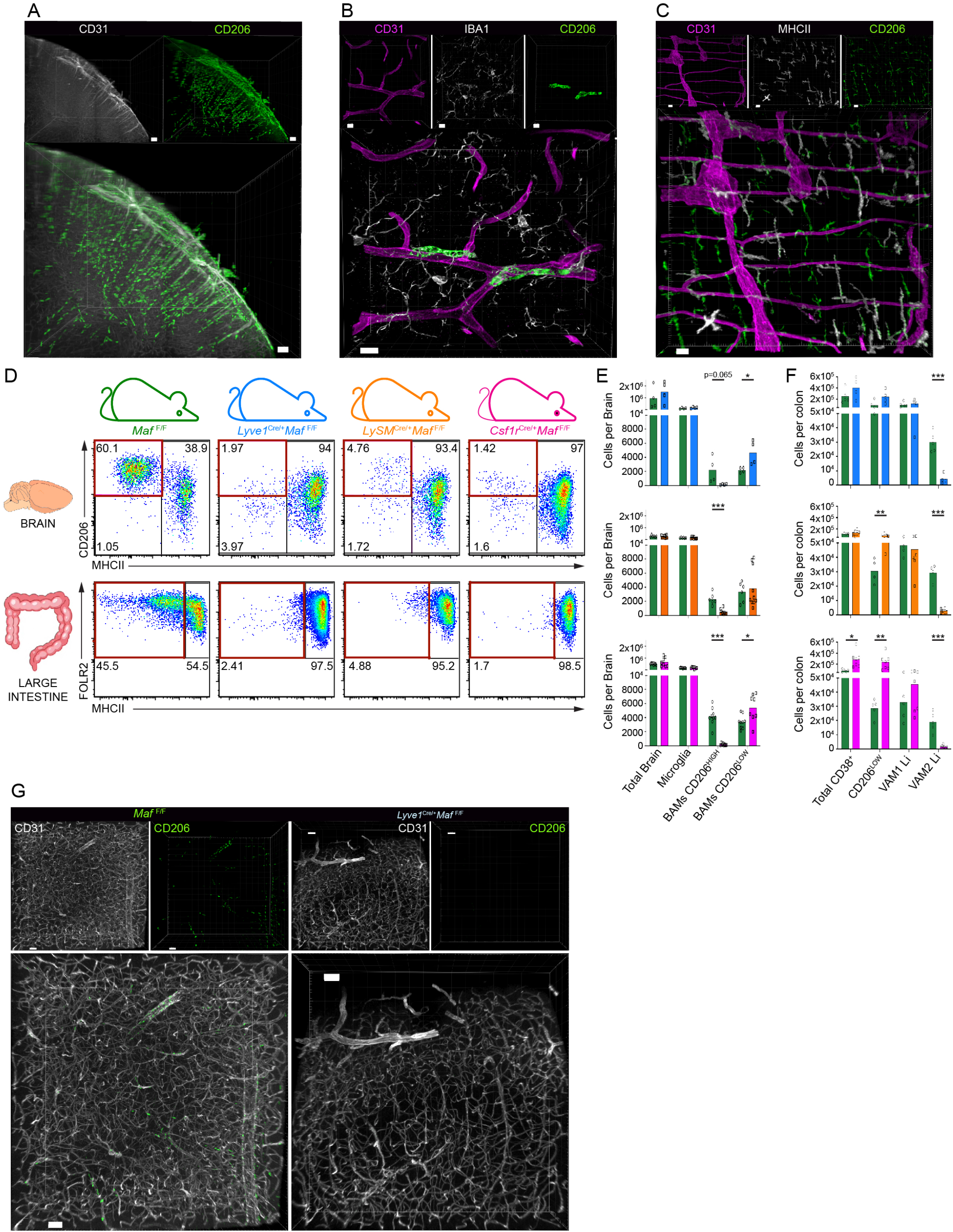
The transcription factor c-MAF is fundamental for the phenotype of VAM like perivascular macrophages in the brain and gut. (A) CD206^HIGH^ macrophages are also tightly associated with blood vessels in the brain and gut of adult WT mice. Lightsheet microscopy of a full mouse brain frontal cortex after clarification and stained with anti-CD206 (Green) and anti-CD31 (White). Scale Bars, 100 μm. 4x magnification. (B) Clarified brain stained with anti-CD206 (Green), anti-CD31 (Magenta) and anti-IBA1 (White). Scale Bars, 10 μm. 63x magnification. (C) Colon *muscularis* stained with anti-CD206 (Green), anti-CD31 (Magenta) and anti-MHCII (White). Scale Bars, 20 μm. 20x magnification. Images representative of at least n = 3. (D) Impact of *Maf* ablation in brain and gut CD206^HIGH^ macrophages. Each column represents macrophages populations of the animal depicted in the top. Each row represents cells from the organ shown in the left. The gates in maroon color highlights the lost of cells with a typical CD206^HIGH^ perivascular phenotype in the conditional knockout models used. Percentage of cells is shown. Representative dot plots of at least n=3. (E) Total brain cell numbers in the different conditional knockout models used in (D). (F) Total colon cell numbers in the different conditional knockout models used in (D). Graph bar colors correspond to the models listed in (D). For each model, littermate controls were used as *Maf*^F/F^ WT controls. (G) Confocal image of a clarified brain of *Lyve1*^Cre^*Maf*^F/F^ and littermate control mice demonstrating the ablation of BAMs CD206^HIGH^. Brains were stained with anti-CD206 (Green), anti-CD31 (white). Scale Bars, 50 μm. 10x magnification. Representative images. n=3. Each dot in the bar graphs represents one animal. Bar graphs display mean values. See also Fig. S3 and S4.

In the brain, non-microglial macrophages have been collectively referred to as Border-Associated Macrophages (BAMs)(*23, 48*) and represent approximately 10% of the all brain macrophages (microglia and non-microglial macrophages)(*49*). We found that adult brain BAMs can be divided into CD38^HIGH^MHC class II^LOW^CD206^HIGH^ (BAMs CD206^HIGH^) and CD38^INT/LOW^MHC class II^HIGH^CD206^LOW^ (BAMs CD206^LOW^) macrophages (Fig. S3A, B). These shared with VAMs high expression levels of LYVE1, Folr2, CD38, TIM4 and CD206 and a lower expression of CD45 (Fig. S3B, C). Light-sheet microscopy of cleared brains revealed that CD206^+^ BAMs were distributed along vessels in the brain cortex, following the vasculature from the meninges into the parenchyma (Fig. 3A). BAMs display clear differences in morphology as well as localization in relation to microglia (Fig. 3B); while microglia display their classical ramified morphology; BAMs are elongated, with an extended surface contacting blood vessels (Fig. 3B).

The intestine contains macrophages that are associated with enteric nervous system plexuses as well as vasculature(*50–53*). We found that the large intestine harbors cells that resemble VAMs in the eWAT. Cells similar to VAM2 CD38^+^Folr2^HIGH^CD206^HIGH^MHCII^LOW^ (VAM2 Li) and VAM1 CD38^+^Folr2^HIGH^CD206^HIGH^MHCII^HIGH^ (VAM1 Li) were clearly distinguishable among the macrophage populations (Fig. S4A, B). The CD206^HIGH^ populations (VAM2 Li and VAM1 Li) comprise around 50% of the CD38^+^ macrophages in the large intestine (Fig. S4A, B) and are mainly located in tight association with blood vessels (Fig. 3C) and around enteric-associated neurons (EANs) (Fig. S4C, D). We observed a similar CD206^HIGH^ population in the ileum muscularis externa layer as well (Fig. S4E); however only the VAM1 related phenotype could be observed in the small intestine (Data not shown). Besides sharing flow cytometry markers and morphological features, the CD206^HIGH^ macrophages in the Brain and Large intestine also display a high endocytic capacity for blood-borne macromolecules (Fig. S3D and fig. S4F), although with kinetic differences between the different organs.

The high similarity shared by all perivascular macrophages in all organs analyzed together with the recent demonstration that BAMs CD206^HIGH^ express high levels of *Maf*(*23*) led us to verify whether the *Maf* dependence observed in the eWAT is also present in other tissues. Similar to what we observed in eWAT, *Maf* gene deletion with all three Cre drivers caused a profound reduction in numbers of BAMs CD206^HIGH^ in the adult brain (Fig. 3D). In this case however, we did not find the emergence of a distinct compensatory population. Using *Csf1r*^Cre^, which is expressed in yolk sac EMP, we could assess the effect in brains of neonatal mice (P5). Indeed, BAM CD206^HIGH^ macrophages were completely absent, without any detectable compensation of the missing macrophages by BAMs CD206^LOW^ (Fig. S3E). Thus, *Maf* gene deletion causes an early ablation of BAMs CD206^HIGH^ in the brain. As animals aged with the continuous *Maf*-dependent ablation of BAMs CD206^HIGH^, monocytes began to enter the brain. While the number of BAMs CD206^HIGH^ remained always exceedingly low, there was an expansion in BAMs CD206^LOW^ macrophages that gradually increased in proportion and number (Fig. 3E). As expected by their low *Maf* expression, microglia numbers were unaffected at all ages (Fig. 3E and fig. S3E). Other myeloid cell types in eWAT and brain were not consistently affected by all three Cre drivers of *Maf*^F/F^, although there were some Cre-specific changes of unclear significance (Fig. S3F). Finally, the impact of *Maf* gene deletion on brain BAMs CD206^HIGH^ can be observed in whole brain cleared images (Fig. 3G and fig. S3G).

In the large intestine, *Maf* deletion also resulted in profound changes in macrophage populations. With all three Cre drivers, as observed in the eWAT, the VAM2 Li disappeared (Fig. 3D and fig. 3F) and a significant increase in the numbers of CD206^LOW^ macrophages is observed (Fig. 3f). The numbers of VAM1 Li and CD38^-^ macrophages were mostly unaffected (Fig. 3F and fig. S4G). Finally, we observed that all CD206^HIGH^FOLR2^+^ macrophages were absent in the small intestine muscularis (Fig. S4H). These data indicate that, in both large and small intestine, c-MAF plays a fundamental role controlling the CD206^HIGH^ macrophages populations of perivascular macrophages. Overall, these analyses establish a crucial role for c-MAF in the establishment and transcriptional programing of perivascular macrophages across tissues.

### *Maf* ablation in perivascular macrophages increases the eWAT vascular network

We next studied in detail the phenotype caused by *Maf* deletion in eWAT VAMs. Because VAMs are juxtaposed to blood vessels we first evaluated the eWAT vascular network. *Maf* ablation with all three Cre drivers resulted in augmented vascular branching in the eWAT, as manifest by increased blood “vessel surface area” (Fig. 4A, B). The increase in vascular branching was mostly observed in the capillaries in eWAT pads and could be detected in young 5 week-old mice (Fig. 4C). Comprehensive morphological characterization indicated that the increased vascular surface area was unlikely due to an overall reduced adipocyte size or any other eWAT gross abnormality (Fig. S5A, B). Despite the alterations in the eWAT vasculature and in VAM gene expression upon *Maf* ablation, the capacity of eWAT VAMs to rapidly uptake blood-borne macromolecules was preserved (Fig. 4D). To assess whether the *Maf* gene itself was responsible for excessive vascular branching in the eWAT, or whether increased branching was the result of the altered VAM phenotype, we crossed *Lyve1*^Cre^ with *Csf1r*^F/F^ mice. The resulting mice had the *Maf* gene unaltered, and VAMs (that are LYVE1^+^) lacked the important gene *Csf1r*(*54, 55*), which is not expressed by LEC. VAMs were reduced in *Lyve1*^Cre^*Csf1r*^F/F^ mice by approximately 2 fold (VAM2) and 3 fold (VAM1) (Fig. 4E). However, the over-representation of Tim4^HIGH^MHCII^HIGH^ VAMs, so prominent in *Maf* gene deleted VAMs (*see* Fig. 1E, F), did not occur in *Lyve1*^Cre^*Csf1r*^F/F^ mice, supporting a specific role for *Maf* in this effect (Fig. 4E). Nevertheless, *Lyve1*^Cre^*Csf1r*^F/F^ mice also had increased vascular branching (Fig. 4F, G), indicating that different genetic perturbations in the VAM compartment can impact eWAT blood vessel architecture. Moreover, the fact that these phenotypes were observed in four complementary conditional knockout models that share VAMs as the only common target (Fig. 1D) strengthens the possibility of a VAM-specific function in regulating vascular branching in the eWAT.

**Fig. 4.**
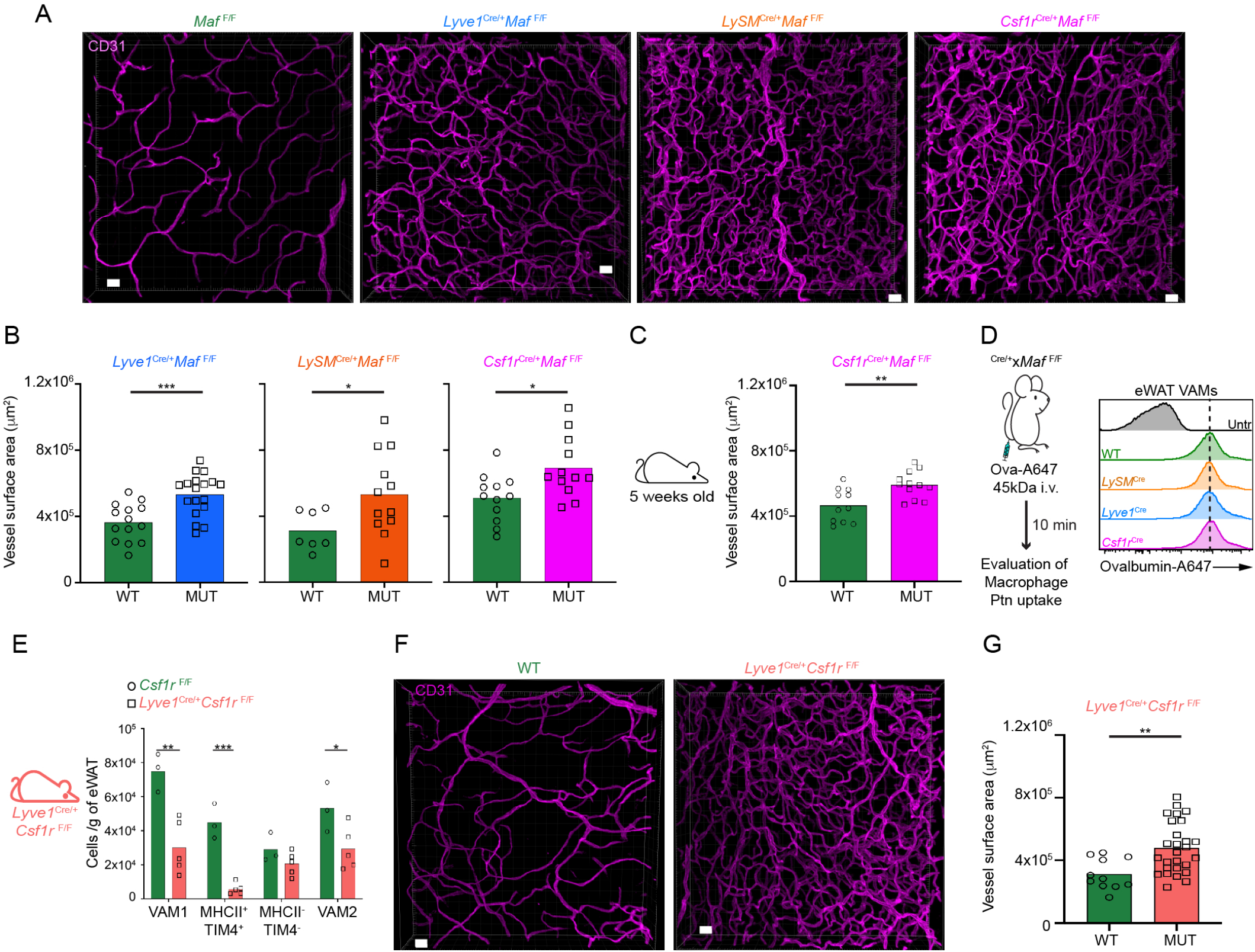
*Maf* ablation in perivascular macrophages leads to increased vascular density in the eWAT. (A) Confocal images of clarified eWAT from animals *Lyve1*^Cre^*Maf*^F/F^, *LySM*^Cre^*Maf*^F/F^, *Csf1r*^Cre^*Maf*^F/F^ and littermate control stained with anti-CD31 (Magenta). Scale Bars, 20 μm. 20x magnification. Representative images. n≥3. (B) Quantification of the vascular surface area assessed by the anti-CD31 staining observed in (A) Three to four 50μm deep images were taken per animal and the vascular surface area reconstructed using the IMARIS software. Each dot represents the quantification for each picture. n≥3. Representative of 3 independent experiments. (C) Quantification of the vascular surface area assessed by the anti-CD31 staining of young 5 weeks old *Csf1r*^Cre^*Maf*^F/F^ mice. Three to four 50μm deep images were taken per animal and the vascular surface area reconstructed using the IMARIS software. Each dot represents the quantification for each picture. n=3. (D) Evaluation of the endocytic capacity of eWAT VAMs in the different animal models used in (A) Ovalbumin-A647 (Ova-A647) was injected i.v. and after the depicted time the uptake of ovalbumin was measured by flow cytometry in the eWAT. Representative histograms; n = 3. The dashed line represents the median fluorescence intensity of the WT VAMs. (E) Distribution of eWAT macrophages per gram of adipose tissue in *Lyve1*^Cre^*Csf1r*^F/F^ mice. Littermate controls were used as *Csf1r*^F/F^ WT controls. Each dot in the bar graphs represents one animal. Representative of 3 independent experiments. (F) *Lyve1*^Cre^*Csf1r*^F/F^ mice display increased vascular density in the eWAT. Confocal images of clarified eWAT stained with anti-CD31 (Magenta). Scale Bars, 20 μm. 20x magnification. Representative images. n≥3. (G) Quantification of the vascular surface area assessed by the anti-CD31 staining observed in (F). Three to four 50μm deep images were taken per animal and the vascular surface area reconstructed using the IMARIS software. Each dot represents the quantification for each picture. n≥3. Representative of 2 independent experiments. Bar graphs display mean values. See also Fig. S5.

### *Maf* deletion in macrophages protects mice from HFD-induced metabolic syndrome

Perivascular macrophages from *Maf*-deficient mice displayed up regulation of important metabolic genes such as *Apoe*, *Hmox* and *Hif3a* that could exert a role in lipid and oxygen metabolism (*see* Fig. 1G, H). Several reports have indicated that metabolic syndrome in a hypercaloric setting develops concomitantly with increasing levels of circulating cholesterol (high LDL), triglycerides and with an increase in adipocyte hypertrophy and hypoxia(*56, 57*).Hence, we opted to test their susceptibility to the metabolic syndrome by feeding them with HFD (Fig. 5A). After 11 weeks of HFD WT littermate controls showed higher fasting glucose serum levels and began started to develop glucose intolerance assessed by glucose tolerance test (GTT), while *Csf1r*^Cre^*Maf*^F/F^ mice were protected from it, not showing any significative difference in relation to Normal Diet (ND) fed counterparts (Fig. 5B, C). The GTT was performed after a 16 hours fast, a time point that gathers information about hepatic glucose metabolism and insulin sensitivity (*58*). Moreover, while ND-fed *Csf1r*^Cre^*Maf*^F/F^ mice gained weight similarly to WT counterparts (Fig. 5D, E), HFD-fed *Csf1r*^Cre^*Maf*^F/F^ mice gained less weight than HFD-fed littermates, and their fat pads were smaller (Fig. 5D, E). Consistently with weight and GTT data, serum triglycerides, which were elevated in littermates fed HFD, were not increased in HFD-fed *Csf1r*^Cre^*Maf*^F/F^ mice (Fig. S6A, B). Additionally, Pyruvate Tolerance Test (PTT) did not reveal differences in gluconeogenesis between *Csf1r*^Cre^*Maf*^F/F^ and littermate controls (Fig. S6C). Furthermore, the two groups displayed similar increase in the circulating triglyceride levels upon intragastric olive oil gavage (Fig. S6D, E), suggesting that the improvements in metabolic parameters observed in *Csf1r*^Cre^*Maf*^F/F^ mice is not due to abnormalities in the hepatic glucose metabolism or intestinal lipid uptake.

**Fig. 5.**
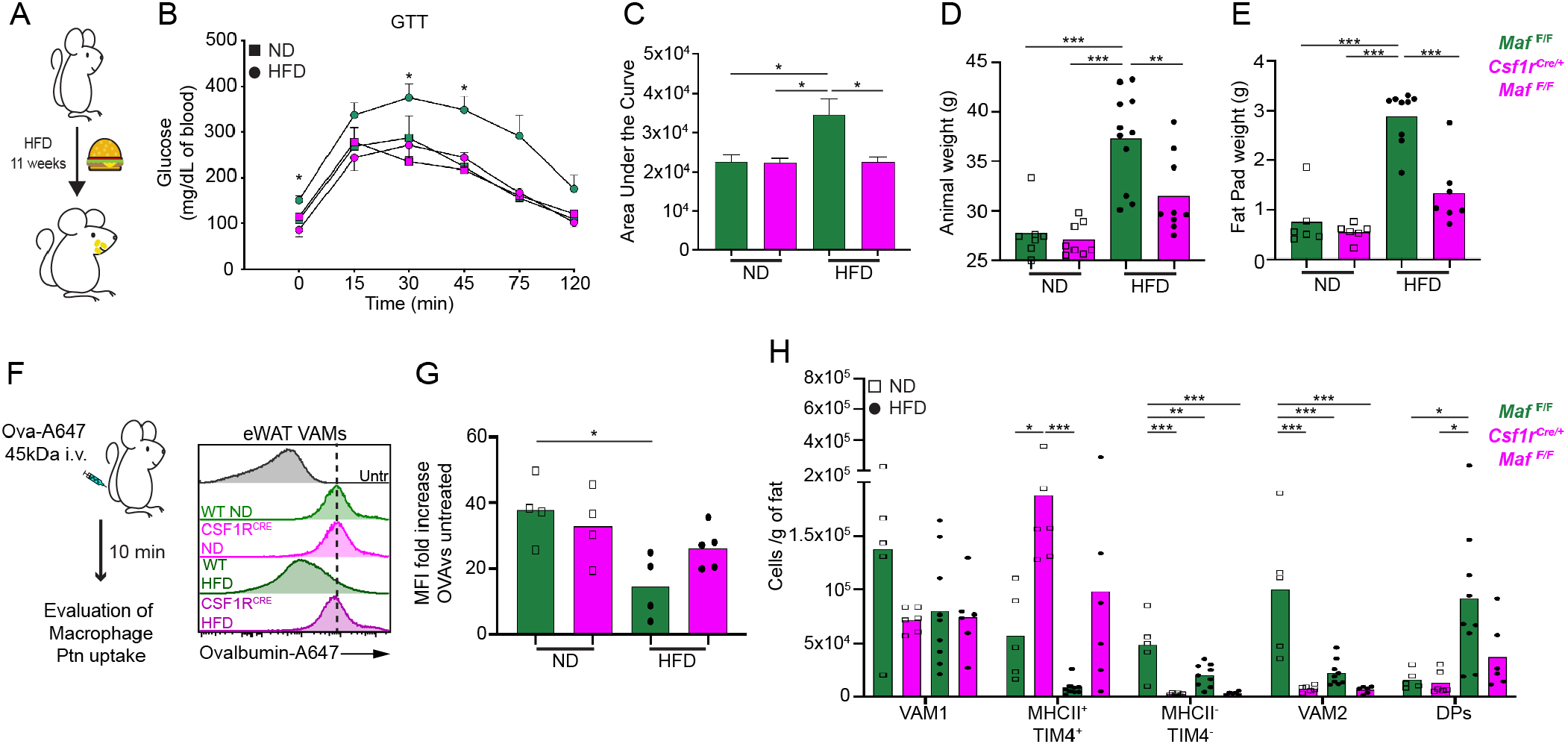
*Maf* ablation in macrophages protects animals from the early onset of metabolic syndrome. (A) Experimental design for (B-H). (B) Glucose tolerance test (GTT) of *Csf1r*^Cre^*Maf*^F/F^ mice and littermate controls under normal or high fat diet (ND or HFD). n>3. Representative of 3 independent experiments. (C) Area under the curve of the GTT displayed in (B) (Arbitrary units). Bar graphs display mean±SEM. (D, E) Animal weight and eWAT fat pads weight of animals depicted in (B). (F) Evaluation of eWAT VAMs endocytic capacity in animals submitted to HFD. Ovalbumin-A647 (Ova-A647) was injected i.v. and after the depicted time the uptake of ovalbumin was measured by flow cytometry in the eWAT. Representative histograms; n ≥4. The dashed line represents the median fluorescence intensity of macrophages from animals under ND regimen. (G) Fold increase of the median ovalbumin-A647 fluorescence intensity after injection as depicted in (F) normalized to the individual autofluorescence of each subpopulation, n≥4. Each dot in the bar graphs represents one animal. Bar graphs display mean values. (H) Distribution of VAMs and DPs (CD11c^+^CD64^+^) macrophages per gram of eWAT in the animals depicted in (B). Each dot in the bar graphs represents one animal. Bar graphs display mean values. DPs: Double positive macrophages were gated as CD45^+^Lin^-^ CD11b^+^CD206^LOW/INT^MHCII^+^CD64^+^CD11c^+^.See also Fig. S6.

Our previous studies showed that VAMs in HFD-fed WT mice have diminished endocytic capacity compared to littermates fed normal diet(*15*). This capacity was also rescued in *Maf*-deficient VAMs (Fig. 5F, G). At the cellular level, the most striking change in the adipose tissue macrophages related to HFD feeding in WT mice was the dramatic increase of BM derived CD11c^+^ DP macrophages(*12, 15*), which constitute only a minor fraction of macrophages in mice fed normal diet. This population did not increase in *Csf1r*^Cre^*Maf*^F/F^ mice upon HFD (Fig. 5H). Furthermore, other myeloid cells that have numbers highly increased in HFD-fed mice, such as Ly6C^HIGH^ monocytes (Mon), Ly6C^LOW^ monocytes and CD11b^-^ dendritic cells, were also found at normal levels in HFD-fed *Csf1r*^Cre^*Maf*^F/F^ mice (Fig. S6F). In conclusion, induction of obesity by HFD, cellular parameters of adipose tissue inflammation, and systemic markers of metabolic syndrome were prevented by ablation of the *Maf* gene in *Csf1r* expressing cells. To our knowledge this is the first study that demonstrates that embryonically derived VAMs can contribute to the pathogenesis of metabolic syndrome. These results define a major role for c-MAF in the transcriptional programing and function of perivascular macrophages; additionally, they reveal a novel role for these cells in the metabolic syndrome.

## DISCUSSION

We found that c-MAF is an essential regulator of embryonically derived perivascular macrophages identity in different organs *in vivo*. Key transcription factors have been identified previously in other macrophage populations; for example c-Fos was shown to regulate the osteoclast lineage(*59*), Gata-6 to regulate peritoneal cavity macrophages(*60–62*), and Spi-C for red pulp macrophages(*63*). The present study defines an essential role for c-MAF in the normal transcriptional program and function of perivascular macrophage *in vivo*, also revealing useful genetic approaches that allow for their examination.

c-MAF is part of the of large Maf family of transcription factors that also include MAFA, MAFB, and Nrl (*29, 64–68*). Among them only MAFB and c-MAF have been reported to be expressed by myeloid cells ((*24, 25, 69–71*) and Immgen). Despite their significant homology, there is little redundancy between the activity of MAFB and c-MAF in myeloid cells. MAFB directly controls the epigenetic landscape of self-renewing genes, limiting the proliferation rate of myeloid cells (*69, 72*). In addition, it was observed that MAFB limits the ability of M-CSF to differentiate hematopoietic stem cells towards the myeloid program (*70*). MAFB can also impact the function of macrophages in several pathological models (*73–76*). In the context of metabolic disease, animals harboring MAFB-deficient macrophages were shown to be more susceptible to obesity in a hyper caloric setting (*75*), an opposite phenotype from our model targeting c-MAF in macrophages under high fat diet. Indeed, others and we reported distinct outcomes in the macrophages transcriptional landscape upon ablation of c-MAF versus MAFB (*27, 30*), however, further studies are necessary to unveil the gene regulatory networks directly associated with these distinct outcomes in pathological models. Finally, while MAFB ablation seems to not impact the macrophages cell identity (*69, 70, 72–76*), we were able to show for the first time *in vivo* that c-MAF defines the identity of embryonically derived perivascular macrophages, such as VAM2.

Macrophages adapt to the tissue environment in which they reside, and are, therefore, different in every tissue(*4, 43, 77*). Nevertheless, we showed that perivascular macrophages in several tissues and organs share key phenotypic and functional properties. The relationship between macrophages and vascular development has been previously addressed. During programmed vascular regression in the developing retina, macrophages induce apoptosis of the endothelial cells and clear the resulting debris(*78, 79*). In the regression of the pupillary vasculature, it was reported that macrophages directly engulf membranes of endothelial cells(*80*). In the fetal testis, macrophages mediate vascular reorganization, driving pruning and angiogenesis(*81*). Our data suggest a direct participation of VAMs in the control of eWAT vasculature development, revealing yet another fundamental functional activity ascribed to macrophages in animal organogenesis(*2–7, 82*). In addition, the increase in the eWAT vascular density upon ablation of c-MAF in VAMs may be associated with the protection from early signs of metabolic syndrome as recently shown (*83*). By identifying c-MAF as a key transcription factor of perivascular macrophages, we provide a genetic target to carry out further mechanistic studies into the role of this macrophage subpopulation in the establishment and maintenance of eWAT normal vasculature.

The striking improvement in obesity and metabolic parameters with manipulation of *Maf* levels in macrophages indicates an important role for tissue resident, embryonically derived, perivascular macrophages in the pathogenesis of obesity-related disease. Historically, the development of diet induced metabolic syndrome is ascribed to the remarkable increase in numbers of bone marrow derived CD11c^+^ (DPs) macrophages within the eWAT (*12, 84*). To determine whether HFD-induced insulin resistance was caused by CD11c^+^ macrophages, CD11c-DTR (diphtheria toxin receptor) was used to ablate all CD11c^+^ cells, including fat DP macrophages, after diphtheria toxin (DT) administration. This treatment resulted in a reduction of HFD-caused WAT macrophage numbers and crown-like structures, and importantly, also caused an improvement of glucose and insulin sensitivity(*85*). However, some caveats of these experiments have been appreciated: 1) CD11c-DTR mice ablate many more cell types than CD11c^+^ macrophages upon DT administration, affecting both innate and adaptive immunity; 2) The authors also showed that DT had a beneficial impact on mice that did not express the DTR, an “off target” impact that has now been recognized in these mice; 3) As we showed, VAMs (not DP) have a remarkable capacity to endocytose blood-borne macromolecules, including proteins. It is therefore possible that some of the off-target events are caused by the DT taken up by VAMs, following death without the need of a DTR.

Our data suggests that the *Maf*-dependent VAMs contribute to the massive inflow of monocyte-derived CD11c^+^ DP macrophages into eWAT of HFD-fed mice, as well as the key glucose tolerance parameters. We previously showed that during chronic HFD feeding, WT VAMs had a significant impairment in their endocytic capacity, and their typical morphology curling around blood vessels, was lost(*15*). In contrast, *Maf* deficiency preserved the high endocytic capacity of VAMs, even during HFD feeding. It is thus possible that the abnormal VAMs in WT mice are the cause of the CD11c^+^ DP infiltration, which contributes to metabolic syndrome. Determining the molecular mechanisms deployed by eWAT VAMs to this phenotype, and if they contribute directly or indirectly to it remains a hurdle due to the intrinsic difficulty to manipulate macrophage subtypes in an organ specific manner. In light of our findings, further development of tools to specifically manipulate subpopulations of macrophages may be crucial to refining our understanding about the contribution of VAMs to the vasculature architecture and systemic metabolism, which may in turn further the establishment of new therapeutic approaches to treat metabolic syndrome.

## MATERIALS AND METHODS

### Mice

C57BL/6J (000664), *Lyve1*^Cre^ (012601), *LySM*^Cre^ (04781), *Csf1r*^Cre^ (021024), *Csf1r*^FLOX^ (021212) and CCR2^RFP^ (017586) strains were purchased from Jackson laboratories. C57BL/6 CD45.1 (564) animals were purchased from Charles River. *Cx3cr1*^CRE-ERT2-YFP^ x *Rosa26*^LSL-DsRed^ was a kind gift of Wenbiao Gan and Dan Littman Lab (*44*). *Maf*^FLOX^ strain was previously described(*34*) and kindly provided by C. Birchmeier. Adult mice at age of 12-16 weeks old were used in the study unless otherwise stated in the Figures. High-fat diet (HFD)-fed male mice were treated with a HFD for at least 11 weeks starting at 4-6 weeks of age. Mice were fed standard chow providing 17% calories from fat (LabDiet Formulab 5008) or a HFD providing 60% calories from fat (Research Diets, D12492i). Glucose tolerance test was performed as described before(*58*). Briefly, animals were submitted to 16hr fasting. After that, 1g/Kg of glucose was injected i.p. and blood glucose levels were measured as depicted in the figures. Time 0 is right before glucose injection. Blood glucose was determined with a One Touch Basic glucometer (Lifescan, Milipitas, California). Pyruvate tolerance test was performed after fasting animals for 16hr. One gram/Kg of sodium pyruvate (Sigma-Aldrich, P5280) was injected i.p. and blood glucose levels were measured as depicted in the figure. Time 0 is right before pyruvate injection. Blood glucose was determined as described above. Mice were sacrificed by CO2 euthanasia and serum was collected when necessary. Serum analyses (Expanded tox panel #60514) were conducted by IDEXX Bioanalytics (North Grafton, MA). Whenever indicated, mice were treated i.v. with 1mg/kg of OVA-A647 (Thermofisher, O34784). For timing of embryonic development, mice were crossed at night, the following day a positive vaginal plug was considered as E0.5. At E9.5 recombination was induced by treatment of pregnant females by gavage with a single dose of 2.5 mg tamoxifen (Sigma) and 1.75 mg progesterone (Sigma) to counteract the mixed oestrogen agonist effects of tamoxifen, which can result in fetal abortions. Animals were housed at NYU Medical Center Animal Facility under SPF conditions. All procedures were approved by New York University School of Medicine Institutional Animal Care and Use Committee.

### Human Visceral Adipose tissue Sections

Anonymous Human visceral adipose tissue was obtained from individuals submitted to bariatric surgery at NYU Langone Tisch Hospital. The samples were considered surgical waste during the procedure. Samples were processed as described below for flow cytometry or fixed overnight at 4°C with 4%PFA for imaging purposes.

### Adipose tissue stromal vascular fraction (SVF) purification

To isolate leukocytes from adipose tissue mice were anesthetized with a mixture containing 12.5mg/mL ketamine, 2.5mg/mL xylazine, and 25 mg/mL acepromazine and intracardially perfused with PBS with 5 mM EDTA. After dissection, the eWAT was minced and incubated for 35 min at 37°C under gentle agitation (150rpm) in cRPMI (2% BSA FFA-free [Fisher BP9704100], 1mM of Sodium Pyruvate [Gibco, 11360-070], Glutamax 1x [Gibco, 35050-061], MEM non-essential amino acids [Gibco, 11140-050], Hepes [Gibco, 15630-080], Pen Strep [Gibco, 15140-122] and RPMI [Gibco, 21870-076]) containing 1mg/mL of collagenase type VIII (Sigma-Aldrich, C2139) and 100μg/mL of DNase I (Sigma-Aldrich, 10104159001). After digestion, the suspension was washed with ice-cold cRPMI and spun down. The floating fat layer was removed and the leukocyte-enriched pellet was resuspended in cRPMI and filtered through a 100 μm cell strainer.

### Isolation of mononuclear cells from brain

Mice were perfused as described. Brain was separated from skull cap, minced and digested with PBS 1X containing 5% FBS, 1 mM Hepes and 240 U/mL collagenase D (Sigma-Aldrich) at 37°C for 35 min under gentle agitation (150rpm). Brain digestion was stopped by adding 10 mM EDTA for 5 min at 37°C. Digested brain was then homogenized, filtered through a 70 μM cell strainer (BD Biosciencies) and centrifuged at 1800 rpm for 10 min at 4°C. Next, cell pellet was resuspended in 38% Percoll (Sigma-Aldrich) and centrifuged at 2000 rpm for 30 min at RT. Following centrifugation, supernatant containing myelin was carefully aspirated and the pellet containing mononuclear cells was used for flow cytometry.

### Isolation of mononuclear cells from gut

#### Small intestine

For the small intestine the distal half (jejunum and ileum) was harvested. Firstly, we removed the mesentery and Peyer’s patches. Small intestine was then cut open longitudinally and washed with PBS 1X to remove luminal contents. Epithelial layer was removed by incubating the tissue once with 1M DTT+10mM Hepes in PBS 1X for 10 min and twice with 30mM EDTA+10mM HEPES in PBS 1X for 15 min. All incubations were performed at 37°C in a mild motion (150rpm) and followed by intense shake of the samples. The samples were then washed with PBS 1X, minced and digested with RPMI containing 10% FBS, 0.01 M Hepes, 400 U/mL collagenase D and 0.1 mg/mL DNAse I for 45 min at 37°C. Digested samples were homogenized, filtered through a 100 μm nylon mesh and centrifuged at 2000 rpm for 10 min at 4°C to obtain the mononuclear cells.

#### Large intestine

The whole large intestine was obtained from anal region up to caecum, and then the mesentery and feces were removed. The luminal side of the large intestine was exposed using a polyethylene tube (8 cm) tightly attached in both boundaries of the intestine and then the epithelial layer was removed as described for the small intestine. Next, the large intestine was minced and digested in RPMI containing 2% BSA, 0.01 M Hepes, 0.2 mg/mL collagenase VIII, 0.1 mg/mL DNAse I and 5 μg/mL dispase for 25 min at 37°C. Digested samples were then homogenized, filtered through a 100 μm cell strainer and centrifuged at 2000 rpm for 10 min at 4°C to obtain mononuclear cells.

### Intestine muscularis dissections

Mice were sacrificed by cervical dislocation and the small intestine ileum (1 cm moving proximal from the ileocecal junction, 6 cm length), or colon (1.5 cm moving distal from cecum and 4 cm moving proximal from the rectum, 6 cm length) was removed. For dissection of the muscularis, the intestinal tissue was then placed on a chilled aluminum block with the serosa facing upward. Curved forceps were then used to carefully remove the muscularis from the mucosa in one intact piece.

### Kidney Capsule eWAT transplant

The eWAT of 5-days old donor mice (P5) was harvested and grafted under the kidney capsule of isofluorane anaesthetized 10-week-old congenic recipient mice using a procedure previously described (*86*). One fat lobe was grafted per kidney. Sixty days post transplant, host animals were sacrificed, the engrafted eWAT was recovered and processed individually as described above.

### Intestinal fat absorption assay

Intestinal fat absorption assay was performed as described before (*87*). In brief, 8-14-week-old male mice were fasted for 6 hrs and received 200 μl olive oil by intragastric gavage, with administration of Triton WR1339 (Sigma, T0307; 0.5 g/kg, i.p.), 30 min prior to gavage. Blood from retro-orbital plexus was collected before gavage and at 1, 2, 3, 4 hr after gavage. Serum triglycerides were measured with the Triglyceride Colorimetric Assay Kit (Cayman, #10010303).

### Antibodies and fluorescent conjugates

Antibodies were purchased from <u>BD-biosciences</u>: Anti-Mouse CD45 (clone 30-F11) BUV 395; Anti-Mouse SiglecF (clone E50-2440) APC-R700; Anti-Mouse TIM4 (Clone RMT4-54) BUV395; Streptavidin-BUV395. <u>Thermofisher</u>: Anti-Mouse CD11b (clone M1/70) APC-E780; Anti-Human CD45 (clone 2D1) APC-E780; Anti-Mouse CD249 (Clone 6C3) Biotin; Anti-Mouse LYVE1 (clone ALY7) Biotin; Anti-Mouse TIM4 (Clone RMT4-54) PE and Percp-E710; Donkey Anti-Rabbit A568 (A10042); Ovalbumin-A647, O34784; Bodipy 493/503, D3922; 4’, 6-diamidino-2-phenylindole (DAPI), D3571 or LiveIdead fixable blue, L34961 were used to exclude dead cells. <u>Biolegend</u>: Anti-Mouse CD11c (clone N418) A647, BV421, BV510 and BV650; Anti-Human CD14 (clone HCD14) PE-CY7; Anti-Human CD16 (clone3G8) A700; Anti-Mouse CD19 (clone 1D3) FITC; Anti-Mouse CD31 (clone 390) A488 and BV421; Anti-Mouse CD38 (clone 90) A647; Anti-Mouse CD45 (clone 30-F11) A700; Anti-Human CD45 (clone 2D1) A700; Anti-Mouse CD64 (clone x54-5/7.1) APC, PE, PE/DAZZLE594, BV421 and Biotin; Anti-Human CD64 (clone 10.1) FITC; Anti-Human CD163 (clone GHI/61) BV421; Anti-Mouse CD206 (clone C068C2) A647, PE-CY7 and PEIDAZZLE594; Anti-Human CD206 (clone 15-2) A647; Anti-Mouse CX3CR1 (clone SA011F11) BV650; Anti-Mouse FOLR2 (clone 10/FR2) PE; Anti-Mouse I-A/I-E (clone M5/114.15.2) A488, Percp-Cy5-5, BV421 and BV711; Anti-Mouse Ly6C (clone HK1.4) A488, Percp-Cy5-5, A700 and BV510; Anti-Mouse Ly6G (clone 1A8) FITC; Anti-Mouse Thy1.2 (clone 30-H12) FITC; Anti-Mouse TIM4 (Clone RMT4-54) A647; Streptavidin-BV421; 405225. <u>Abcam</u>: Anti-Mouse LYVE1 (ab14917); Anti-Human CD31 (AB32457); Donkey Anti-Goat A568 (AB175704); <u>R&D</u>: Anti-Mouse CD31 (AF3628); Anti-Mouse LYVE1 (AF2125); Anti-Human LYVE1 (AF2089). <u>Jackson Immunoresearch</u>: Donkey Anti-Rat A647 (712-605-153); Donkey Anti-Rat BV421 (712-675-153), Donkey Anti-Goat A488 (705-545-147); Donkey Anti-Goat A647 (705-605-147); Normal Donkey Serum (017-000-121). <u>WAKO</u>: Anti-Mouse IBA1 (019-19741). Human polyclonal Anti-Hu (ANNA-1) was a kind gift from V. Lennon.

### Flow cytometry and cell sorting

Flow cytometry data were acquired on an LSR-II flow cytometer (Becton Dickinson) and analyzed using FlowJo software (Tree Star 8.7). All stainings were performed as recommended by the antibody manufacturer. All samples were pre incubated with Fc block (2.4G2, BE0307 bioxcell) for 15 min at 4°C. FACS sorting was performed using an ARIA II sorter using a 100μm nozzle (Beckton Dickinson). Flow cytometry of adipose tissue macrophages is made somewhat difficult because of their high autofluorescence. To minimize the impact of autofluorescence, we meticulously selected the fluorochromes for each cell marker, and used fluorescence minus one controls (FMOs).

### Bone marrow chimeras

Recipient mice were lethally irradiated (1000 rads) and reconstituted with bone marrow cells from mice as depicted in the figures. After 2 months, mice were sacrificed, and organs of interest analyzed.

### RNA isolation and sequencing

Total RNA from sorted target cell populations was isolated using TRIzol LS (Invitrogen; 10296010) followed by DNase I (Qiagen, 79254) treatment and cleanup with RNeasy Plus Micro kit (Qiagen, 74034). RNA quality was assessed using pico bioanalyser chips (Agilent, 5067-1513). Only RNAs displaying a RNA integrity number (RIN) of 9 or higher were used for downstream procedures. RNAseq libraries were prepared using the Nugen Ovation Trio low input RNA Library Systems V2 (Nugen; 0507-08) according to the manufacturer’s instructions by the NYU Genome Technology Center. Pooled libraries were sequenced as 50 nucleotide, paired-end reads on an Illumina NextSeq 500 using v4 chemistry.

### RNAseq data quality assessment and visualization

Illumina sequencing adapters and reads with Phred quality scores lower than 20 were removed with Trimmomatic. Trimmed reads were mapped to the *Mus musculus* genome Ensembl annotation release 91 using STAR v2.5.3a with default settings. The number of reads uniquely mapping to each gene feature in the corresponding annotation file was determined using featureCounts. The resulting count tables were passed to R for further analyses.

### Differential expression analysis and motif identification

DESeq2 was used for differential expression analysis. Samples from different experiments performed on different days were analyzed separately. Consistency between replicates was checked using principal component analysis (PCA) and Euclidean distance-based hierarchical clustering on normalized counts. Genes with an average of less than 130 normalized counts across all samples were removed from volcano plots to visualize only the most highly expressed genes. Genes were considered significantly differential when the adjusted P value was less than 0.05.

### Whole-mount intestine muscularis immunofluorescence

Following intestine dissection, the intestine muscularis tissue was pinned down on a Sylgard-coated plate, covered with PBS/4% PFA and fixed overnight at 4°C. Following tissue fixation, samples were washed in DPBS (minimum 4x 15min), followed by permeabilization in 0.5 % Triton X-100/0.05 % Tween-20/4 μg heparin (PTxwH) for a minimum of 2 hr at RT with gentle agitation. Samples were then blocked for 2 hr in blocking buffer (PTxwH with 5% donkey serum) for 2 hr at RT with gentle agitation. Primary antibodies were added to blocking buffer at appropriate concentrations and incubated for 2-3 days at 4°C. Following primary antibody incubation, samples were washed in PTxwH (minimum 4x 15min), followed by incubation with secondary antibody in PTxwH at appropriate concentrations overnight at 4°C. Following secondary antibody incubation, samples were again washed in PTxwH, and mounted with FluoroMount G on slides with 1 ½ coverslips. Slides were kept in the dark at 4°C until they were imaged(*52*).

### Adipo-Clear of brain and eWAT

#### Sample Collection

the method was performed as described before(*35*). Animals were anesthetized as described before and an intracardiac perfusion/fixation was performed with 1x PBS followed by 4% PFA. All harvested samples were post-fixed in 4% PFA at 4°C overnight. Fixed samples were washed in PBS for 1 hr three times at RT.

#### Delipidation and Permeabilization

fixed samples were washed in 20%, 40%, 60%, 80% methanol in H2O/0.1% Triton X-100/0.3 M glycine (B1N buffer, pH 7), and 100% methanol for 1hr each. Sample were then delipidated with 100% dichloromethane (DCM; Sigma-Aldrich) for 1h three times. After delipidation, samples were washed in 100% methanol for 1hr min twice, then in 80%, 60%, 40%, 20% methanol in B1N buffer for 1hr each step. All procedures above were carried out at 4°C with shaking. Samples were then washed in B1N for 30 min twice followed by PBS/0.1% Triton X-100/0.05% Tween 20/2 μg/mL heparin (PTxwH buffer) for 1hr twice before further staining procedures.

#### Immunolabeling

For Immnolabeling of the eWAT we used just the apical region of the epidydimal lobe for staining. For the Brain, whole brain or half hemisphere (Sagital cut) was stained. Samples were incubated in primary antibody dilutions in PTxwH for 4 days at RT. After primary antibody incubation, samples were washed in PTxwH for 5 min, 10 min, 15 min, 30 min, 1 hr, 2 hr, 4 hr, and overnight, and then incubated in secondary antibody dilutions in PTxwH for 4 days at RT. Samples were finally washed in PTwH for 5 min, 10 min, 15 min, 30 min, 1 hr, 2 hr, 4 hr, and overnight.

#### Tissue Clearing

samples were dehydrated in 25%, 50%, 75%, 100%, 100% methanol/H2O series for 30 min at each step at RT. Following dehydration, samples were washed with 100% DCM for 2hr twice, followed by an overnight clearing step in dibenzyl ether (DBE; Sigma-Aldrich). Samples were stored at RT in the dark until imaging.

#### 3D Imaging

Whenever indicated for whole-tissue imaging we used a light-sheet microscope (Ultramicroscope II, LaVision Biotec) equipped with 1.3X (used for whole-tissue views with low-magnification) and 4X objective lenses (used for high-magnification views) and an sCMOs camera (Andor Neo). Images were acquired with the ImspectorPro software (LaVision BioTec). Samples were placed in an imaging reservoir filled with DBE and illuminated from the side by the laser light sheet. The samples were scanned with the 488 nm, 640 nm and 790 nm laser channels and with a step-size of 3 μm for 1.3x objective and 2.5 μm for 4x objective. For high magnification and measurement of eWAT vessel architecture clarified fat pieces and brain were placed in a μ-Slide 2 Well Glass Bottom (Ibidi, #80287) containing DBE as mounting media and imaged at room temperature on an inverted confocal microscope (Zeiss 710 MP; optical lenses 20x N.A. 0.8 and 63x N.A. 1.4). For eWAT vessel architecture, 4 images per tissue/animal were acquired.

#### Image Processing

All whole-tissue images were generated using Imaris x64 software (version 9.2, Bitplane). 3D reconstruction was performed using the “volume rendering” function. The vessel surface area was obtained by using the Batch tool to process all images in an unbiased way. Briefly, all samples were submitted to a median filter and then, using the “*Surfaces*” algorithm as described by the manufacturer, the vascular tree was reconstructed for 50 μm z-stacks (each stack separated by 0.87 μm) and the vessel surface area obtained. Optical sections were generated using the “snapshot” tool.

### Confocal microscopy of neonatal WAT macrophages

White adipose tissue was stained as described(*88*), with modifications. Mice were euthanized and slowly perfused by intracardiac injection with 20 mL of PBS/5mM EDTA. Epididymal fat pads were excised and fixed for 1h in PBS/1% paraformaldehyde with gentle shaking at 4°C. After that, samples were blocked for 1 h in PtxwH buffer (0.5 % Triton X-100/0.05 % Tween-20/4 μg heparin) containing 5% Donkey serum (blocking buffer) with gentle rocking at room temperature. Primary antibodies were diluted in blocking buffer and added to fat samples for 3 days at 4°C. Samples were then washed 6 times with cold PTxwH buffer (30min at 4°C each). After that samples were incubated with secondary antibodies for 3 days at 4°C. The tissue was washed again as described before and incubated with blocking buffer-diluted Bodipy (Thermofisher # D3922) for 1h at room temp. Fat pads were imaged at room temperature on an inverted confocal microscope (Zeiss 710 MP; optical lenses 20x N.A. 0.8 and 63x N.A. 1.4) by placing the pad in Fluoromount-G (Southern Biotech, 0100-01) in a chambered coverslip. Zen software was used for image acquisition. ImageJ software, Fiji version 1.0 or Imaris/Bitplane software were used for contrast, brightness and pseudo-color adjustments.

### Histology

Tissues were fixed in 4% paraformaldehyde and embedded in paraffin. Twenty-micrometre sections were stained with haematoxylin and eosin. Bright-field color images were acquired in a Zeiss Axio Observed using 20x magnification lenses (N.A. 0.8). The adipocyte diameter was measured using the ImageJ software.

### Statistical analysis

Mean, SD and SEM values were calculated with GraphPad Prism (version 8.4.1, GraphPad Software). Error bars represent ± SEM. Unpaired Student’s *t* test was used to compare two variables. For 3 or more variables one way ANOVA was used with a turkey post test, as indicated in each figure legend. P-values < 0.05 were considered significant. Statistics symbols were determined as: * = p<0.05, ** = p<0.01 and *** = p<0.001.

### Data and code availability

Data generated by RNA sequencing are deposited in the NCBI Gene Expression Omnibus (GEO) database and are accessible under GEO: GSE148606. All software used is available online, either freely or from a commercial supplier.

## Acknowledgments

We thank Dr. Maria Lafaille and Dr. Gabriel Victora for valuable comments on the manuscript. We acknowledge NYULH DART Microscopy Lab for assistance with microscopy work. Microscopy shared resource is supported by the Cancer Center Support Grant, P30CA016087. We thank Alireza Khodadadi-Jamayran from NYU Langone Medical Center Applied Bioinformatics Laboratory for initial assembly of processed RNAseq data. We acknowledge NYULH DART Genome Technology Center for assistance with RNA sequencing. This a shared resource partially supported by the Laura and Isaac Perlmutter Cancer Center, Cancer Center Support Grant P30CA016087. J.Z. Kitoko and C.P. Queiroz were supported by fellowships from the Coordination for the Improvement of Higher Education Personnel (CAPES-Brazil). H.M. Silva was supported by a fellowship from the National Council for Scientific and Technological Development (CNPq – Brazil) and a postdoctoral fellowship from Dr. Bernard B. Levine.

## Author Contributions

Conceptualization, H.M.S., D.M., M.T.B. and J.J.L.; Methodology, H.M.S., J.Z.K., C.P.Q., L.K., F.M., K.L.Y and C.R.F.; Software, H.M.S.; Validation, H.M.S., J.Z.K., C.P.Q., L.K., F.M., K.L.Y and C.R.F.; Formal Analysis, H.M.S., J.Z.K., C.P.Q., L.K. and F.M.; Investigation, H.M.S., J.Z.K., C.P.Q., L.K., F.M., K.L.Y and C.R.F.; Data Curation, H.M.S.; Writing – Original Draft, H.M.S and J.J.L.; Writing – Review & Editing, H.M.S., J.Z.K., C.P.Q., L.K., F.M., K.L.Y., C.R.F., D.R.L., M.T.B., D.M. and J.J.L.; Visualization, H.M.S., J.Z.K. and C.P.Q.,; Supervision, H.M.S. and J.J.L.; Project Administration, H.M.S. and J.J.L; Funding Acquisition, J.J.L.

## Materials & Correspondence

Hernandez Moura Silva or Juan J. Lafaille

## SUPPLEMENTARY MATERIALS

Fig. S1 (Related to Main Fig. 1). Distribution of perivascular macrophages expressing CD38^+^Folr2^+^CD206^HIGH^ in the eWAT.

Fig. S2 (Related to Main Fig. 2). Adult VAM2s do not develop from bone marrow progenitors.

Fig. S3 (Related to Main Fig. 3). Brain BAMs CD206^HIGH^ share key similarities and *Maf*-dependence with eWAT VAM2.

Fig. S4 (Related to Main Fig. 3). CD206^HIGH^ macrophages from the large intestine display a phenotype that resembles eWAT VAMs.

Fig. S5 (Related to Main Fig. 4). eWAT adipocyte size distribution is mostly preserved in young *Csf1r*^Cre^*Maf*^F/F^ mice.

Fig. S6 (Related to Main Fig. 5). Animals *Csf1r*^Cre^*Maf*^F/F^ display improved metabolic parameters under HFD.

## SUPPLEMENTARY FIGURE LEGENDS

**Fig. S1.**
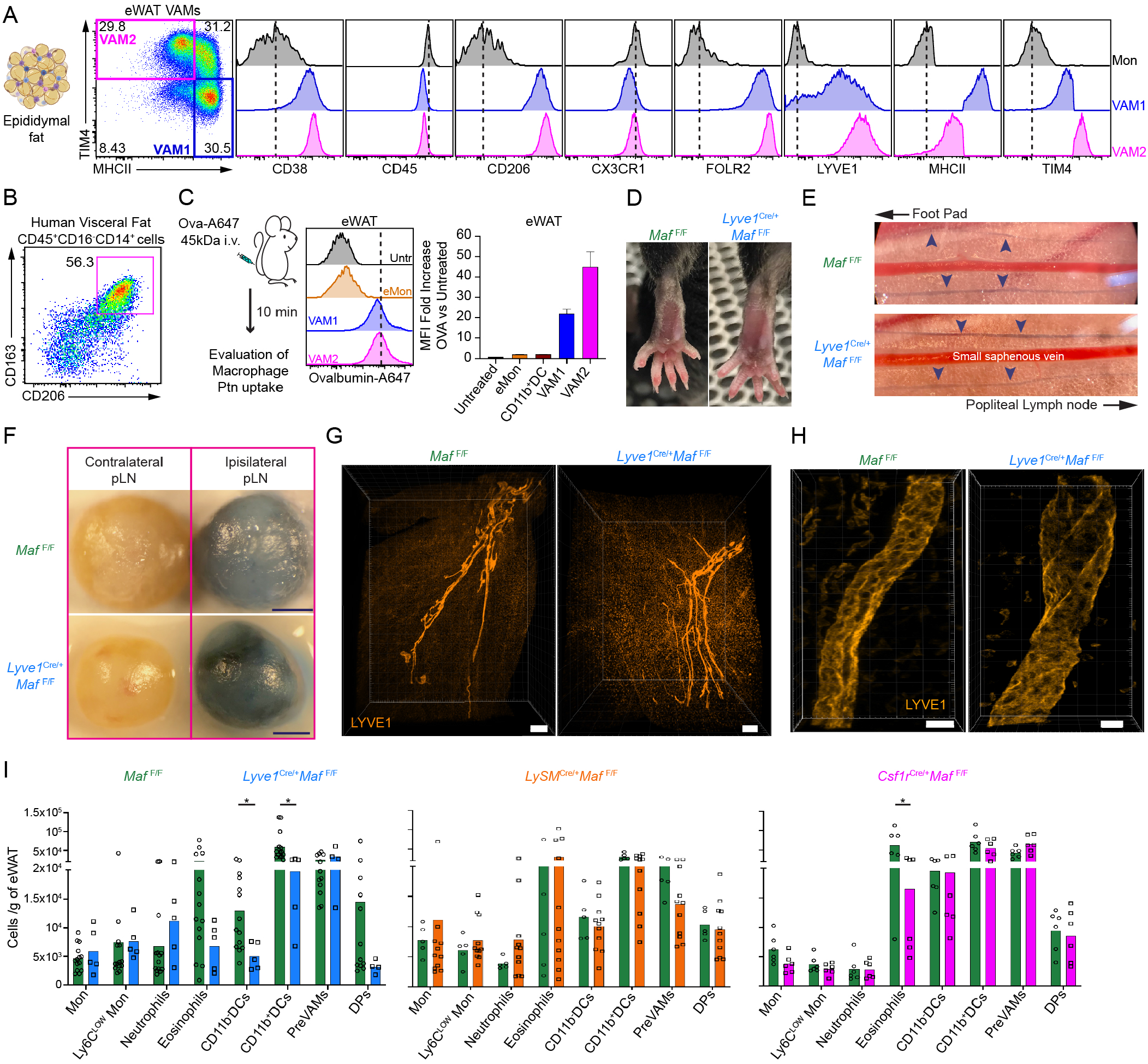
Distribution of perivascular macrophages expressing CD38^+^Folr2^+^CD206^HIGH^ in the eWAT. (A) Flow cytometry analysis of cell surface markers expressed by VAMs in adult WT C57BL/6 mice. eWAT VAMs were gated as CD45^+^Lin^-^CD11b^+^CD64^+^CD206^HIGH^. The magenta and blue gates correspond to VAM2 and VAM1, respectively. (B) Flow cytometry analysis of human visceral fat macrophages. Percentage of cells is shown. Representative dot plot. n=3. (C) VAMs are highly endocytic *in vivo*. Ovalbumin-A647 (Ova-A647) was injected i.v. and after the depicted time the uptake of Ovalbumin was measured by flow cytometry. VAMs gated as mentioned in (A). Representative histograms; n = 3. Dashed lines represent the median fluorescence intensity of the VAM2. Bar graphs show the fold increase of the median ovalbumin-A647 fluorescence intensity after injection normalized to the individual autofluorescence of each subpopulation, n = 3. CD11b^+^DCs gated as CD45^+^Lin^-^CD11b^+^CD64^-^CD11c^+^MHCII^+^. Bar graphs display mean±SEM. (D) *Lyve1*^Cre^*Maf*^F/F^ mice do not show abnormality in liquid drainage from extremities. Foot pad were not swollen in these animals. Representative figure n=5. (E) *Lyve1*^Cre^*Maf*^F/F^ mice display functional lymphatic vessels. Evans blue dye (40mg/ml) was injected in one hind foot pad of *Lyve1 ^Cre^Maf*^F/F^ or littermate controls and the dye drainage through lymphatic vessels around the small saphenous vein was visualized 30 min post injection. Blue arrowheads are indicating evans blue loaded lymphatic vessels. Representative figure. n=3 for each group. (F) Evans blue dye is efficiently drained into the Popliteal lymph node of *Lyve1*^Cre^*Maf*^F/F^. Lymph nodes of the experiment depicted in (E) are shown. pLN = Popliteal lymph node. Scale Bars, 500 μm. Representative figure. n=3 for each group. (G) *Lyve1*^Cre^*Maf*^F/F^ mice display a preserved capacity to form a lymphatic vessel tree within the eWAT. Whole eWAT fat pad were clarified, stained with LYVE1 (Orange) and imaged using a light sheet microscope. Scale Bars, 300 μm. 4x magnification. (H) Lymphatic vessels of *Lyve1*^Cre^*Maf*^F/F^ display a similar structure to those observed in littermate controls. Clarified eWAT were stained with LYVE1 (Orange) and imaged using a confocal microscope. Scale Bars, 20 μm. 20x magnification. Representative images. (I) Distribution of myeloid cells per gram of eWAT in the different conditional knockout models used in main Fig. 1E. Graph bar colors correspond to the models listed in main Fig. 1E. For each model, littermate controls were used as *Maf*^F/F^ WT controls. Ly6C^LOW^ Mon: Ly6C^LOW^ Monocytes were gated as CD45^+^Lin^-^CD11b^+^CX3CR1^+^MHCII^-^ Ly6C^LOW^. Neutrophils were gated as CD45^+^CD11b^+^MHCII^-^Ly6G^+^. Eosinophils were gated as CD45^+^CD11b^+^MHCII^-^SSC^HIGH^SiglecF^+^. PreVAMs were gated as CD45^+^Lin^-^ CD11b^+^CD206^LOW/INT^MHCII^+^CD64^+^CD11c^-^. DPs: Double positive macrophages were gated as CD45^+^Lin^-^ CD11b^+^CD206^LOW/INT^MHCII^+^CD64^+^CD11c^+^. CD11b^-^DCs were gated as CD45^+^Lin^-^ CD11b^-^CD64^-^CD11c^+^MHCII^+^. Ptn: Protein. Lin: CD90^+^CD19^+^Ly6G^+^SiglecF^+^.

**Fig. S2.**
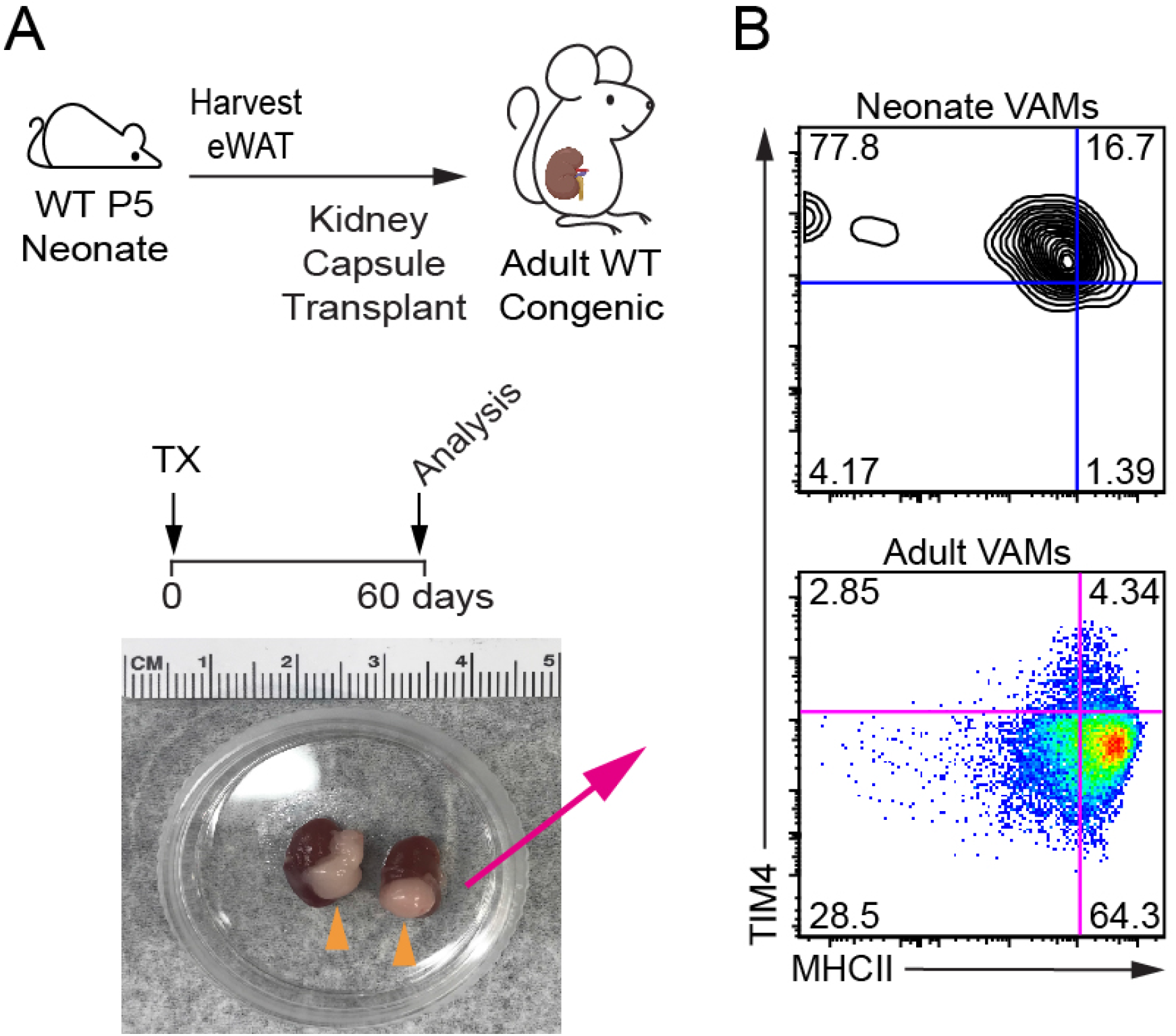
Adult VAM2s do not develop from bone marrow progenitors. (A) Experimental design for (B). (B) Phenotype of organ resident VAMs (Neonate) *versus* BM derived VAMs (Adult VAMs) originated from the transplant recipient that migrated to the transplanted eWAT under the kidney capsule as explained in (A) VAMs (gated as CD45^+^Lin^-^ CD11b^+^CD64^+^CD206^HIGH^). Representative dot plot. n=4. Percentage of cells is shown.

**Fig. S3.**
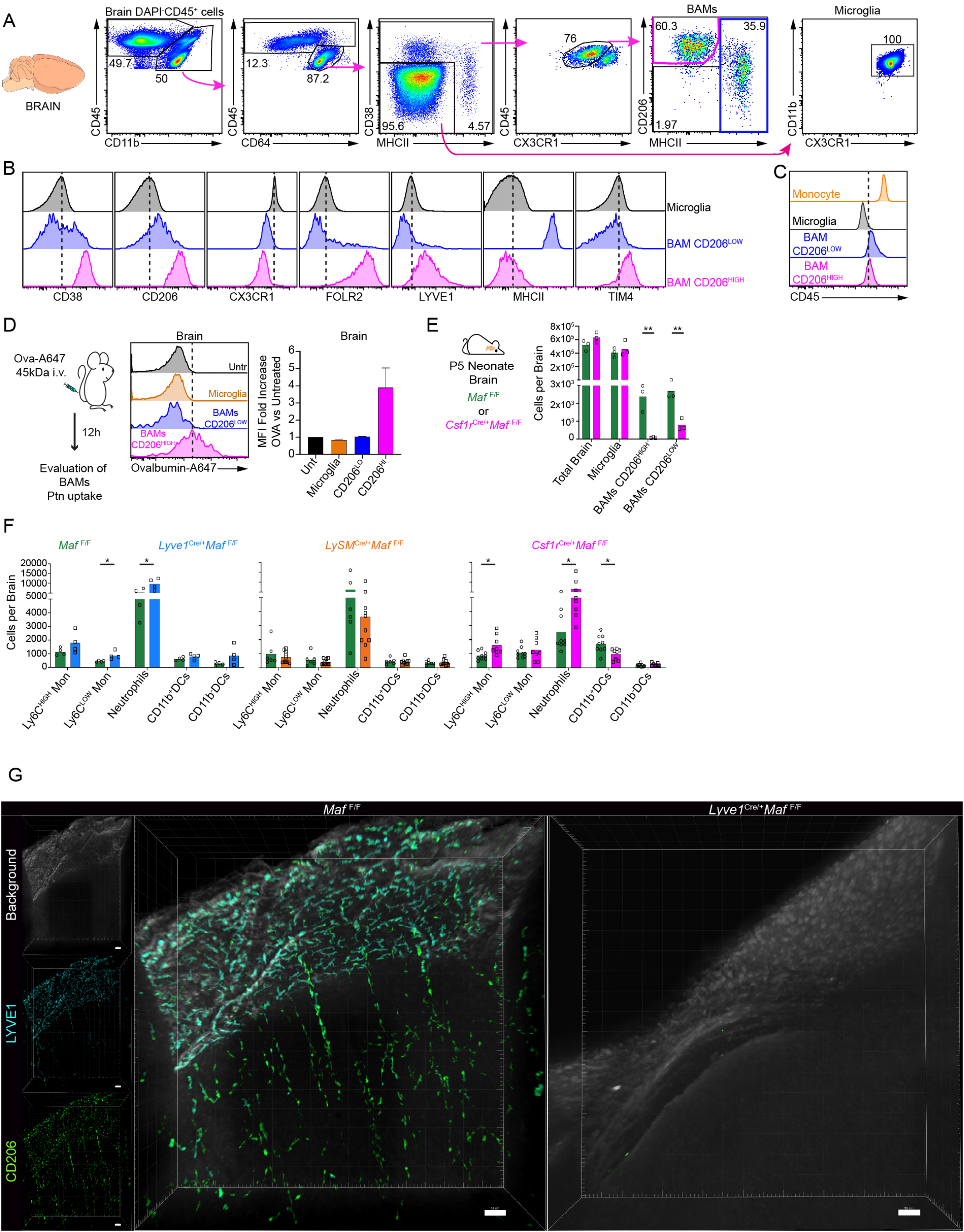
Brain BAMs CD206^HIGH^ share key similarities and *Maf*-dependence with eWAT VAM2. (A) Flow cytometry gate strategy used to analyze brain perivascular macrophages. Percentage of cells is shown. (B) Flow cytometry analysis of cell surface markers expressed by BAMs. Histograms represent the pink and blue gates depicted in the BAMs dot plot in (A). Representative histograms of at least n=5. Dashed lines represent the median fluorescence intensity of microglia as a comparative measure in the organ. (C) BAMs CD206^HIGH^ express lower levels of CD45 in relation to bone marrow derived monocytes. n≥5. Monocytes were gated as CD45^+^Lin^-^CD11b^+^CX3CR1^+^MHCII^-^Ly6C^HIGH^. (D) BAMs CD206^HIGH^ are highly endocytic *in vivo*. Ovalbumin-A647 (Ova-A647) was injected i.v. and after the depicted time the uptake of Ovalbumin was measured by flow cytometry. BAMs gated as mentioned in (A). Representative histograms; n = 3. Dashed lines represent the median fluorescence intensity of the BAMs CD206^HIGH^. Bar graphs show the fold increase of the median ovalbumin-A647 fluorescence intensity after injection normalized to the individual autofluorescence of each subpopulation, n = 3. Bar graphs display mean±SEM. (E) Neonate Brains (P5) show early ablation of BAMs CD206^HIGH^. Total Brain cell numbers in *Csf1r*^Cre^*Maf*^F/F^. Littermate controls were used as *Maf*^F/F^ WT controls. n=3. Representative of 2 independent experiments. (F) Distribution of myeloid cells per brain in the different conditional knockout models used in main Fig. 3 D, E. For each model, littermate controls were used as *Maf*^F/F^ WT controls. (G) Confocal image of clarified brains of *Lyve1*^Cre^*Maf*^F/F^ and littermate control mice demonstrating the ablation of BAMs CD206^HIGH^. Brains were stained with anti-CD206 (Green), anti-LYVE1 (Cyan) and the background (White). Scale Bars, 20 μm. 20x magnification. Representative images. n=3. Ly6C^LOW^ Mon: Ly6C^LOW^ Monocytes were gated as CD45^+^Lin^-^CD11b^+^CX3CR1^+^MHCII^-^Ly6C^LOW^. Neutrophils were gated as CD45^+^CD11b^+^MHCII^-^Ly6G^+^. CD11b^-^DCs were gated as CD45^+^Lin^-^CD11b^-^CD64^-^ CD11c^+^MHCII^+^. CD11b^+^DCs were gated as CD45^+^Lin^-^CD11b^+^CD64^-^CD11c^+^MHCII^+^. Ptn: Protein. Lin: CD90^+^CD19^+^Ly6G^+^SiglecF^+^.

**Fig. S4.**
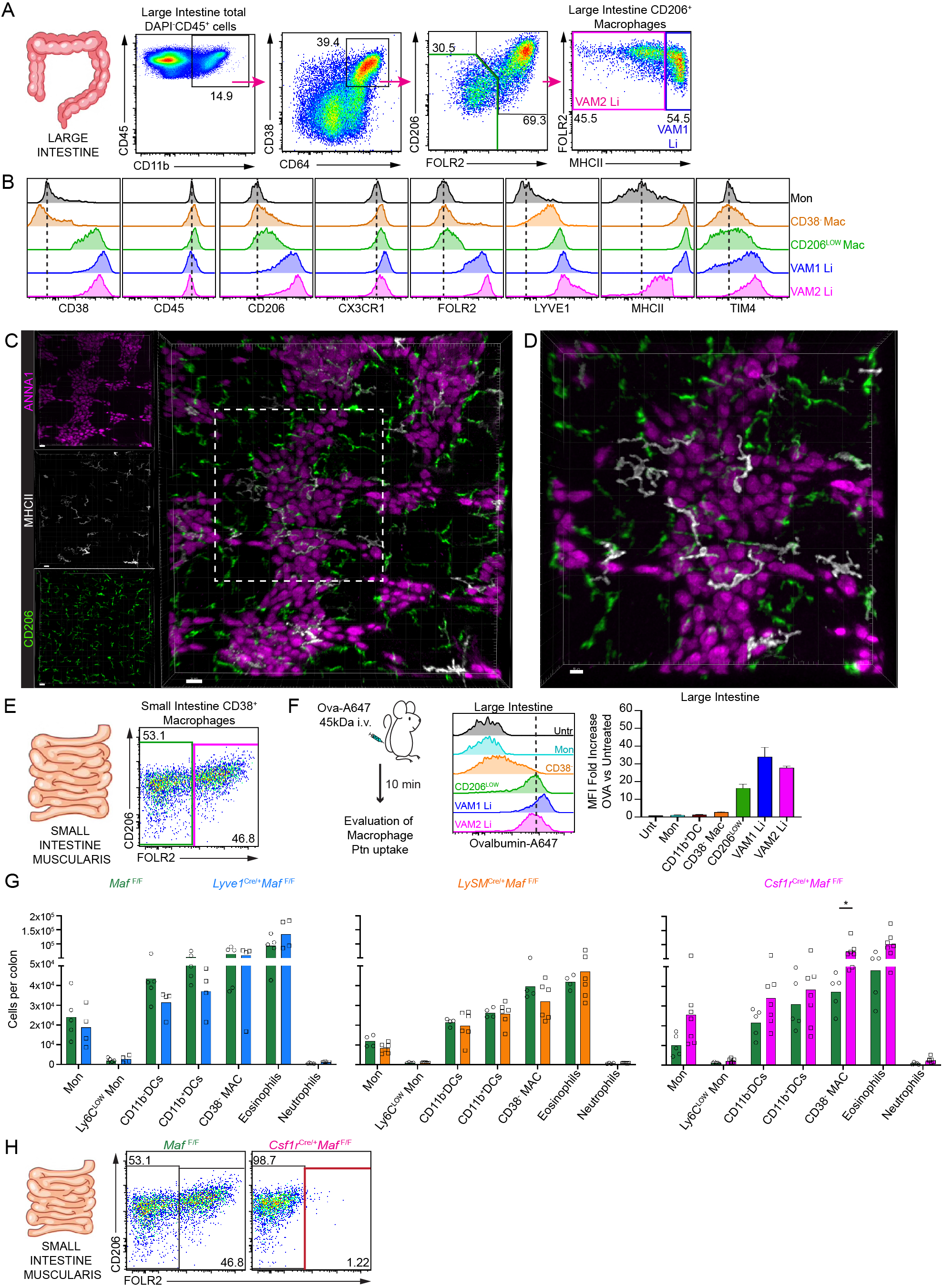
CD206^HIGH^ macrophages from the large intestine display a phenotype that resembles eWAT VAMs. (A) Flow cytometry gate strategy used to analyze large intestine (Li) macrophages of mice used in this study. Percentage of cells is shown. (B) Flow cytometry analysis of cell surface markers expressed by VAMs from Li. Histograms represent the magenta, blue and green gates depicted in the Li macrophages in (A). CD38^-^ Mac: CD38^-^ macrophages were gate as CD45^+^CD11b^+^CD64^+^CD38^-^MHCII^+^. Mon: Monocytes were gated as CD45^+^Lin^-^ CD11b^+^CX3CR1^+^MHCII^-^Ly6C^HIGH^. Representative histograms of at least n=5. Dashed lines represent the median fluorescence intensity of monocytes as a comparative measure in the organ. (C) VAMs Li are also localized in close proximity to enteric-associated neurons in the large intestine myenteric plexus of adult WT mice. Colon stained with anti-CD206 (Green), anti-ANNA-1 (Magenta – neuron body) and anti-MHCII (White). Scale Bars, 20 μm. 20x magnification. n = 3. (D) Same as in **c** in higher details. Figure depicts the white box shown in (C) showing in details the close proximity of perivascular macrophages to enteric-associated neurons. Scale Bars, 10 μm. 20x magnification. (E) The small intestine *muscularis* layer also display VAM related macrophages (pink) and non-related (green) perivascular macrophages. Cells were gated as depicted in (A). Percentage of cells is shown. n=3. (F) VAMs Li are highly endocytic *in vivo*. Ovalbumin-A647 (Ova-A647) was injected i.v. and after the depicted time the uptake of Ovalbumin was measured by flow cytometry. VAMs Li gated as mentioned in (A). Representative histograms; n = 3. Dashed lines represent the median fluorescence intensity of the VAM2 Li. Bar graphs show the fold increase of the median ovalbumin-A647 fluorescence intensity after injection normalized to the individual autofluorescence of each subpopulation, n = 3. CD11b^+^DCs gated as CD45^+^Lin^-^CD11b^+^CD64^-^CD11c^+^MHCII^+^. Bar graphs display mean±SEM. (G) Distribution of myeloid cells per colon in the different conditional knockout models used in main fig. 3D and fig. 3F. For each model, littermate controls were used as *Maf* ^F/F^ WT controls. (H) Representative dot plot demonstrating the impact of *Maf* deletion in the small intestine *muscularis* layer of animals *Csf1r*^Cre^*Maf*^F/F^. Each dot in the bar graphs represents one animal. Bar graphs display mean values. Ly6C^LOW^ Mon: Ly6C^LOW^ Monocytes were gated as CD45^+^Lin^-^CD11b^+^CX3CR1^+^MHCII^-^Ly6C^LOW^. Neutrophils were gated as CD45^+^CD11b^+^MHCII^-^ Ly6G^+^. Eosinophils were gated as CD45^+^CD11b^+^MHCII^-^SSC^HIGH^SiglecF^+^. CD11b^-^DCs were gated as CD45^+^Lin^-^CD11b^-^CD64^-^CD11c^+^MHCII^+^. CD11b^+^DCs were gated as CD45^+^Lin^-^ CD11b^+^CD64^-^CD11c^+^MHCII^+^. Mac: Macrophage. Ptn: Protein. Lin: CD90^+^CD19^+^Ly6G^+^SiglecF^+^.

**Fig. S5.**
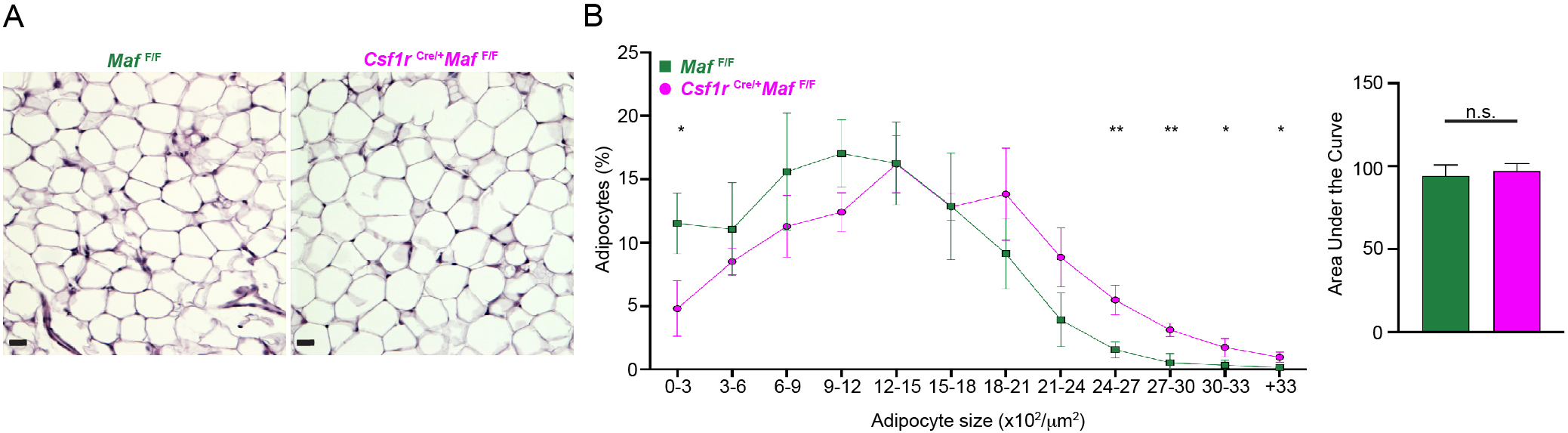
eWAT adipocyte size distribution is mostly preserved in young *Csf1r*^Cre^*Maf*^F/F^ mice. (A) eWAT h&e stainingIsectioning of the animals depicted in the main fig. 4C. Scale Bars, 20 μm. 20x magnification. Representative images. n=3. (B) Distribution of adipocytes sizes over the total number of adipocytes evaluated from animals depicted in (A). For each animal at least 6 randomly acquired h&e sections were analyzed. An average of 380 adipocytes were evaluated per animal using ImageJ software. Area under the curve was evaluated using the graph prism v.8 software.

**Fig. S6.**
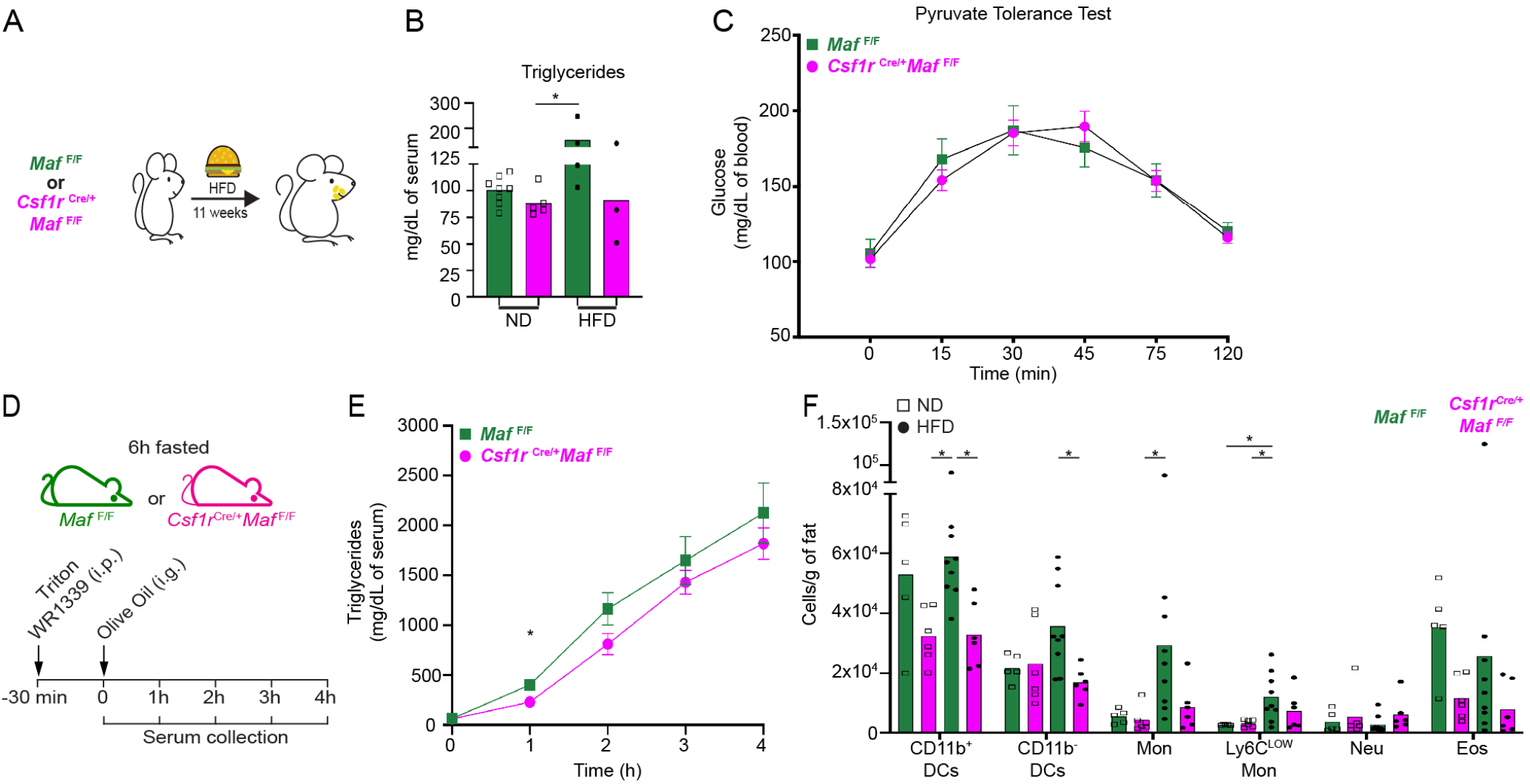
Animals *Csf1r*^Cre^*Maf*^F/F^ display improved metabolic parameters under HFD. (A) Experimental design for (B and F). (B) Triglycerides levels obtained from animals depicted in main fig. 5 B. n≥3. (C) Pyruvate tolerance test to evaluate liver gluconeogenesis. ND-fed *Csf1r*^Cre^*Maf*^F/F^ mice and littermate controls were fasted for 16 hours. Sodium pyruvate (1g/kg) was then injected intraperitoneally and glucose blood levels were evaluated in the depicted times. n = 8 to 10 per group. (D) Experimental design for (E). (E) Serum triglyceride content in ND-fed *Csf1r*^Cre^*Maf*^F/F^ mice and littermate controls after gavage with 200 μl of olive oil. Mice were fasted for 6 hours before gavage. Triton WR1339 (0.5 g/kg) was injected intraperitoneally 30 min before gavage. n=8, per group. (F) Distribution of myeloid cells per gram of eWAT in *Csf1r*^Cre^*Maf*^F/F^ mice submitted to HFD used in main fig. 5 B. eMon: eWAT Ly6C^HIGH^ Monocytes. Ly6C^LOW^ Mon: Ly6C^LOW^ Monocytes. Neu: Neutrophils. Eos: Eosinophils. DCs: Dendritic cells. n≥3. Representative of 3 independent experiments. Each dot in the bar graphs represents one animal. Bar graphs display mean values.

